# Small Molecule Inhibitors Targeting Methyltransferase-Like (METTL) Proteins Against Hepatocellular Carcinoma: A Comprehensive Drug Repurposing Approach

**DOI:** 10.1101/2023.03.11.532187

**Authors:** Md Niaz Morshed, Md Sorwer Alam Parvez, Rakibul Islam Akanda, Manash Kumar Saha, Jannatul Fardous, Mohammad Jakir Hosen

**Affiliations:** Department of Genetic Engineering & Biotechnology, Shahjalal University of Science & Technology, Sylhet 3114, Bangladesh; Department of Drug Discovery Medicine, Kyoto University Graduate School of Medicine, Sakyo ward, 606-8507 Kyoto, Japan; Chittagong Medical College, Chittagong University, Chittagong, Bangladesh

**Keywords:** Methyltransferase inhibitors, hepatocellular carcinoma, drug repurposing, multi-target drug design

## Abstract

An efficient and durable multi-targeted therapeutic drug against hepatocellular carcinoma (HCC) has recently been a growing concern for tackling the chemoresistance of approved anti-HCC drugs. Recent studies indicated that methyltransferase-like (METTL) proteins including METTL1, METTL3, METTL6, METTL16, and METTL18, have overexpressed and associated with the progression of HCC malignancy, and making them excellent biomarkers. Here, we present a series of bioinformatics study including novel compound repurposing approach, molecular docking, pharmacophore modeling, and molecular dynamic simulation, which revealed two first-in-class highly potent catalytic multi-target inhibitors (ZINC70666503 and ZINC13000658 with 87% and 82% drug scores, respectively) of methyltransferase-like proteins. Comparatively, these two inhibitors showed a notable binding affinity against studied METTL proteins. Furthermore, ADME and toxicity analysis suggested that these two commercially available compounds have good drug-likeliness properties with no potent toxic effects. Of note, the molecular dynamics study supported their conformational stability and high selectivity at the pocket of proteins’ adenosine moiety of S-Adenosyl Methionine. However, this comprehensive analysis needs *in vivo* validation to facilitate multi-targeting therapeutic development against hepatocellular carcinoma.

## 1. INTRODUCTION

Being the fourth leading cause of cancer-related deaths, hepatocellular carcinoma (HCC) is one of the deadliest human malignancies worldwide. Infection of Hepatitis B and C virus,, diabetes, dietary toxins, heavy alcohol intake, and nonalcoholic fatty liver disease are the major risk factors for HCC [1-3]. As the molecular mechanism of hepatocellular carcinoma initiation and progression are elusive, it limits the treatment strategies for patients with advanced HCC [4]. However, in recent years, researchers identified several intracellular molecules as biomarkers to detect HCC progression and develop anti-HCC drugs [5,6]. Until now, Food and Drug Administration (FDA) has approved various anti-cancer agents for treating HCC, such as Sorafenib and Lenvatinib [7]. Nevertheless, these drugs or other classical chemotherapeutic agents, tyrosine kinase inhibitors, and novel immune-sensitizing strategies of HCC treatment failed to exert an efficient and durable response toward HCC [8,9].

Recent studies have shown that several members of the Methyltransferase-like (METTL) protein family could play a vital role in HCC and various human cancer [10]. The METTL protein family has a structurally conserved methyltransferase domain which contains an S-Adenosyl Methionine (SAM) binding domain that transfers methyl groups to nucleic acids, proteins, lipids, and small molecules. Among this family, Methyltransferase like protein 1 (METTL1) plays a role in HCC progression by promoting HCC cell proliferation and migration via suppressing PTEN (Phosphatase and tensin homolog) signaling [11,12]. METTL3 regulates glycolysis activity in HCC cells, and the UBC9/SUMOylated Mettl3/Snail axis mediated HCC progression [13,14]. On the other hand, METTL6 and METTL18 also upregulated in HCC tumor tissues, and their suppression correlates with proliferation, invasion, and migration of HCC cells [15,16]. Moreover, METTL16 is another identified m6A methyltransferase, which negatively correlated with decreased RAB11B-AS1 transcript expression in HCC tissues. Thus, the METTL16–RAB11B-AS1 regulatory axis and these above-mentioned methyltransferase family proteins become novel biomarkers and therapeutic targets for hepatocellular carcinoma [17].

Unfortunately, there are no promising therapeutic strategies to specifically block the catalytically active site of these METTL proteins except STM2457, a selective first-in-class SAM-competitive inhibitor of METTL3 [18]. Thus, it is of concern to identify and develop highly-specific and effective anti-HCC drugs targeting multi methyltransferase-like proteins overexpressed in HCC. Bacteria and viruses also need SAM-dependent RNA methyltransferase for their survival and replication, and several studies identified inhibitors targeting their SAM-binding pocket for developing potential antiviral and antibacterial strategies [19-27]. In addition, studies also searched a few SAM-dependent DNA methyltransferase (DNMT) inhibitors against different human malignancies that might be probable anti-cancer drugs [28-31]. In this study, we considered studying the inhibitory potential of those SAM-dependent microbial RNA methyltransferase inhibitors, human DNMT inhibitors, and all identified METTL3 inhibitors against all HCC-associated methyltransferase-like (METTL) proteins for developing anti-HCC therapeutics; a novel perspective of drug repurposing [32-39].

Computational approaches have evolved for drug prediction based on drug candidate– ligand/receptor interaction that has implied searching for potential small molecule inhibitors with a high binding affinity [40]. In parallel, we established a novel pipeline of multi-targeting computer-aided drug design (CADD) along with a pharmacophore modeling-based approach by using *in silico* state-of-the-art techniques, including molecular docking, virtual screening, ADME/T, and molecular dynamics simulation to find potential therapeutic agents for addressing HCC chemoresistance toward existing anti-HCC therapeutics.

## 2. MATERIALS AND METHODS

### 2.1 Retrieval of METTL proteins and preparation of drug compounds

The crystal structure of the methyltransferase-like protein METTL1 [PDB ID: 7PL1], METTL3-14 complex [PDB ID: 7O2I], METTL6 [PDB ID: 7F1E], METTL16 [PDB ID: 6B91], and

METTL18 [PDB ID: 4RFQ] with the resolution of 1.85 Å, 3.00 Å, 2.59 Å, 1.94 Å, and 2.40 Å, respectively, were retrieved from the RCSB Protein Data Bank (PDB) [41]. Furthermore, the three-dimensional (3D) PDB structures of 39 microbial tRNA methyltransferase inhibitors, 20 DNA methyltransferase inhibitors, and 14 studied METTL3 small molecule inhibitors were retrieved from the PubChem database (S.table 1) [42].

### 2.2 Receptor-based screening of drug compounds and active sites identification

Using AutoDock Vina software for molecular docking approaches, selected compounds were screened against all METTL proteins [43]. At first, we processed the crystal structure of all METTL proteins by separating the ligand molecules, removing the water and other complex molecules, and adding polar hydrogen by Discovery Studio 2021 and AutoDock Vina [44]. After preparing the PDB structures of the drug candidates, the highest binding affinity and interactive amino acids were assessed by exposing them to all five proteins. The grid boxes with box shape-size (x, y, z) and shape-center (x, y, z) were set for METTL1, METTL3, METTL6, METTL16, and METTL18 (S.table 2), according to the proteins’ binding sites taken from UniProtKB for molecular docking [45]. PyMOL was further used to analyze all protein-ligand interactions and visualize the interaction sites [46]. Finally, based on the highest binding energy, we selected the top compounds for each protein and analyzed the protein-ligand complexes to identify interacted residues by Protein-Ligand Interaction Profiler (PLIP) [47]. Additionally, the non-covalent interactions (hydrogen bonds, water bridges, salt bridges, halogen bonds, hydrophobic interactions, π-stacking, π-cation interactions, and metal complexes) in those protein-ligand complexes were also analyzed by PLIP.

### 2.3 Quantum Mechanical (QM) calculation

The electronic properties of drug molecules are of significant interest in drug designing, which can be effectively studied through Density Functional Theory (DFT) analysis [48]. In this investigation, the DFT calculation was implemented by employing B3LYP (Becke exchange functional, which combined Lee, Yang, and Parrs (LYP) correlation functional) [49]. For single-point DFT analysis by ORCA version 4.2.1, top inhibitors with the highest molecular docking score were treated with B3LYP and RIJCOSX approximation after minimal geometry optimization by Avogadro 1.2.0n [50-52]. In this calculation, we analyzed frontier molecular orbitals, namely highest occupied molecular orbital (HOMO), lowest unoccupied molecular orbital (LUMO), their energy gap difference, and chemical potentials. Then, the frontier energies (ε) of HOMO and LUMO were used to measure the hardness and softness of selected compounds. The hardness, η (eq.1) and softness, S (eq.2) of the drugs were measured by the Parr and Pearson interpretation equation and Koopmans theorem equation [53-54]. From this, we can evaluate an atom’s ability to receive electrons and higher reactivity, indicating a higher softness value. The following equations measure chemical hardness and softness.

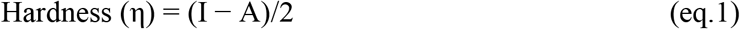

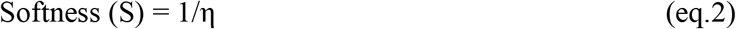

In these equations, ‘I’ refer to the ionization potential (− EHOMO) and ‘A’ denotes the electron affinity (− ELUMO).

### 2.4 Pharmacophore modeling and virtual screening of the ZINC database

Pharmacophore features modeling was done by using PharmaGIST, that are essential for the interaction of inhibitors with METTL proteins [55]. This study used the topmost inhibitors to generate the pharmacophore model. Then, the generated pharmacophore model were imported to ZINCPharmer, and a library of chemical compound was created by screening the ZINC database to carry out further virtual screening [56]. Finally, a virtual screening of the library of compounds against all studied HCC-associated METTL proteins were conducted by AutoDock Vina to find the best drug candidates.

### 2.5 Drug likeness properties analysis of the candidate small molecule inhibitors

Absorption, Distribution, Metabolism, and Excretion (ADME) properties were assessed by the SwissADME server to successfully evaluate the physiochemistry, lipophilicity, water solubility, drug-likeness, and pharmacokinetics properties of potential drug candidates [57]. SwissADME computed the physicochemical descriptors (Formula, Molecular weight, Molar Refractivity, TPSA), lipophilicity (Log Po/w (iLOGP), Log Po/w (XLOGP3), Log Po/w (WLOGP), Log Po/w (MLOGP), Log Po/w (SILICOS-IT), Consensus, Log Po/w), pharmacokinetic parameters (GI absorption, CYP1A2 inhibitor, CYP2C19 inhibitor, CYP2C9 inhibitor, CYP2D6 inhibitor, CYP3A4 inhibitor, Log Kp; skin permeation) and water solubility (Log S: SILICOS-IT, Solubility). Besides, Blood Brain Barrier (BBB) permeability were also computed by adopting the BOILED-Egg yolk method for the screened compounds. Additionally, admetSAR was used to investigate the undesired effects of toxicity, and bioavailability [58]. Finally, drug likeliness and drug score as well as tumorigenicity, mutagenicity, and reproductive effects were assessed by OSIRIS Property Explorer [59].

### 2.6 Evaluation of the docking performance

We further evaluated our AutoDock Vina docking results of final drug candidates against studied proteins using two popular online docking servers, SwissDock and CB-Dock, followed by site-specific re-docking by AutoDock Vina [60,61]. SwissDock can automatically set up the protein and ligand structures and provide convenient visualization and analysis of docking predictions. On the other hand, CB-Dock can automatically predict binding modes and binding sites of a given protein and calculate the centers and sizes with a novel curvature-based cavity detection approach. Thereby, it becomes easier to predict proteins’ binding site cavities where they interact with the inhibitors. For site specific re-docking by AutoDock Vina, the grid box size (x, y, z) and centre (x, y, z) was set as 27 Å × 30 Å × 27 Å and 46 Å × -20 Å × 6 Å; 27 Å × 27 Å × 27 Å and -43 Å × 31 Å × 26 Å; 27 Å × 27 Å × 27 Å and 39 Å × -25 Å × 15 Å; 27 Å × 27 Å × 27 Å and -7 Å × 8 Å × 20 Å; and 27 Å × 27 Å × 27 Å and 6 Å × 8 Å × 28 Å for METTL1, METTL3, METTL6, METTL16, and METTL18, respectively. After re-docking with three different docking tools, we further selected the best conformations of final drug candidates for interaction analysis by PLIP.

### 2.7 Molecular dynamics simulation

Molecular Dynamics (MD) is an important step for evaluating the stability of the receptor-ligand complex. Hence, 100 ns MD simulation was employed to evaluate the binding stability of the final two compounds to all studied METTL proteins at their active site cavity. GROMACS software suit was used for the simulation with GROMOS96 43a1 forced field for 100ns [62]. We used the PRODRG web server for the preparation of the ligand and generation of ligand topology [63]. The 5000 steps steepest descent method was used for the minimization process and SPC was selected as water model for correct density and dielectric permittivity of water. The approximate number of frame per simulation was set to 1000 and NVT/NPT equilibration system was operated under 300K temperature and 1 bar pressure. Additionally, during molecular dynamics simulation study, the ligand-receptor root mean square deviation (RMSD), RMS fluctuation, number of hydrogen bonds, and radius of gyration (Rg) were also calculated to determine the stability and compactness of protein inhibitor complexes. We further performed the MM/GBSA (molecular mechanics, the generalized born model, and solvent accessibility) analysis by using Linux operating system to calculate the ligand binding free energies and ligand strain energies for docked lead compounds with METTL proteins [64].

## 3. RESULTS

### 3.1. Screening of selected inhibitors against METTL proteins

All of the retrieved 73 inhibitor molecules (S.table 1) were employed and the scoring function of AutoDock Vina was utilized to gain structural and atomistic scale interaction insight involving the binding modes of METTL proteins and the inhibitors. Remarkably, the maximum listed inhibitors showed good binding affinity and lower binding energy than SAM, the catalytic activator of methyltransferase proteins. CID76318201, a DNMT inhibitor, was found to have the highest negative binding energy (−12.6 kcal/mol) when interacting with the METTL6 (Table 1). Moreover, CID135449332 (−10.7 kcal/mol), CID5494506 (−10.7 kcal/mol), CID344265 (−11.5 kcal/mol) were also found to be the topmost METTL1, METTL3, and METTL18 inhibitors with high binding affinities, respectively. The molecular docking results of these 73 compounds against all METTL proteins are included in S.table 3.

**Table 1:**
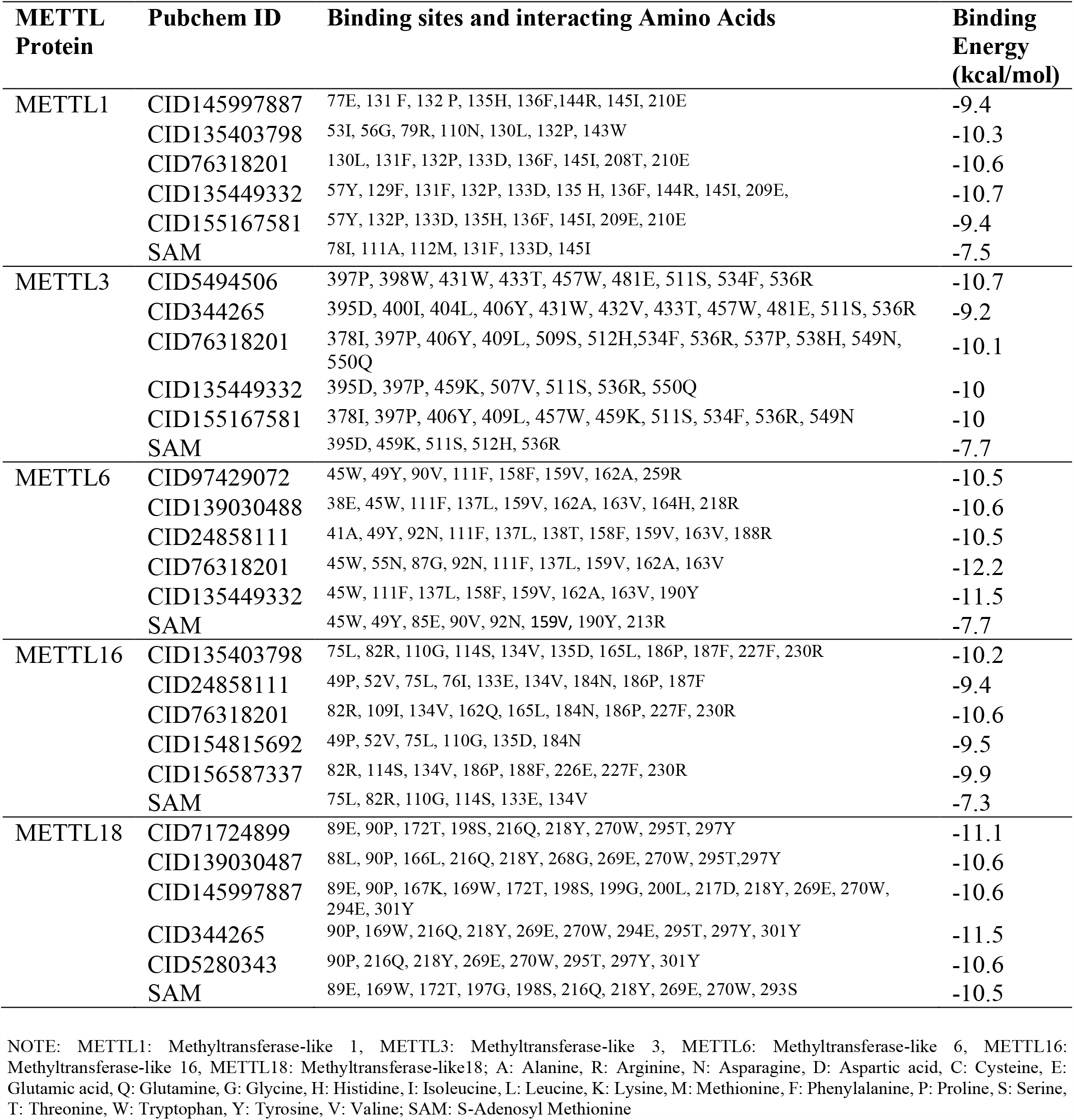
Molecular docking energy and interacted amino acids of hepatocellular carcinoma associated methyltransferase family proteins with top ligands and S-Adenosyl Methionine (SAM)

### 3.2 Structural insights into inhibitor binding hotspots of METTL proteins

The protein-ligand interactions at the docked site of all 73 compounds and five Methyltransferase-like proteins were visualized and analyzed by PyMOL and protein-ligand interaction profiler (PLIP), respectively, to identify proteins’ drug binding active sites. The top five compounds from molecular docking evaluation for each METTL protein were taken to analyze common interactive sites of proteins (Table 1). It was found that compounds with highest binding energy share the same binding modes, forming essential bonds with amino acids residues of 53I, 56G, 57Y, 77E, 79R, 110N, 129F, 130L, 131F, 132P, 133D, 135 H, 136F, 143W, 144R, 145I, 208T, 209E, 210E of METTL1 inside the SAM binding active site which is in amino acid position of 78I, 111A, 112M, 131F, 133D, 145I. For METTL3, there were 23 amino acid positions 378I, 395D, 397P, 398W, 400I, 404L, 406Y, 409L, 431W, 433T, 457W, 459K, 481E, 507V, 509S, 511S, 512H, 534F, 536R, 537P, 538H, 549N, 550Q; which were abundantly found in protein-ligand interaction site (Figure 1). Furthermore, analysis of the METTL6-inhibitors complexes showed that the binding of the ligands was stabilized through the non-covalent bonds with the active site residues, mostly, 45W, 111F, 137L, 159V, 162A, 163V (Table 1). Most surprisingly, the binding patterns analysis indicated that all of the selected top compounds interacted at the binding pocket of the adenosine moiety of SAM. Additionally, the binding pattern of top five compounds for each METTL proteins were highly similar and they shared common residues when interacting. The binding modes of energetically top five inhibitors inside the active sites of respective METTL proteins are depicted in Figure 1 and S.figure 1, and the interacting binding site residues are given in the Table 1. As seen from data listed in S.table 3, 12 inhibitors showed average docking scores > -9.0 kcal/mol.

**Figure 1.**
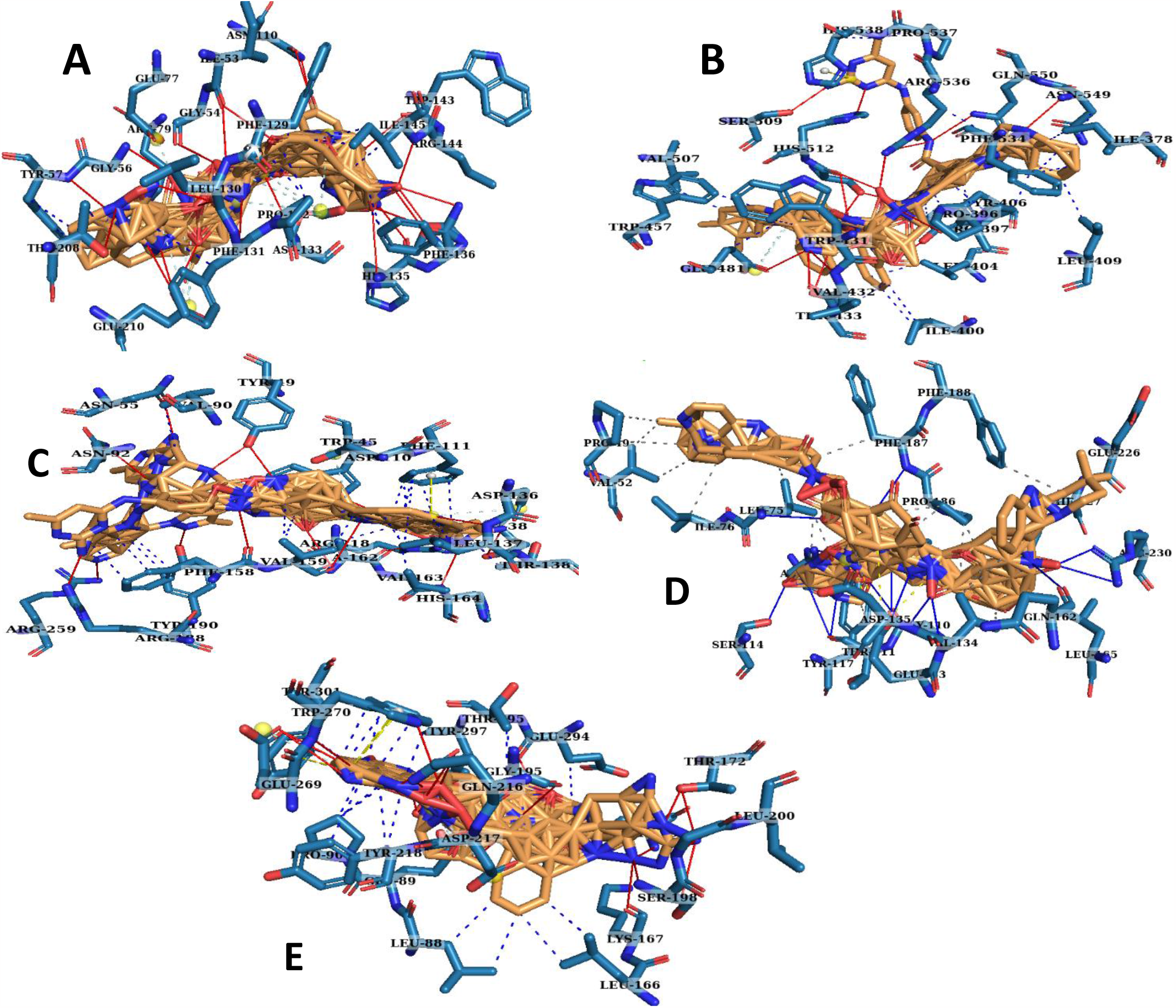
Structural analysis of active sites of METTL family proteins and interacting amino acids at the active sites. Compounds specifically bound at the active site cavity of METTL1, and METTL3, METTL6, METTL16, and METTL18 by interacting with the amino acids shown in A, B, C, D, and E, respectively. Here at A, B, C, D, and E, superimposed top 5 drug compounds in each category are Lemon and interacted amino acids are in Sky blue lines.

### 3.3 Quantum Mechanical Analysis

Compounds which showed high average binding energy were further subjected to DFT calculations, particularly frontier molecular orbitals energy. In the DFT calculations, we calculated the hardness and softness values of 12 energetically top inhibitors (S.table 4). Among them, five compounds showed the highest softness value (>0.6 eV) compared to others. CID139030487, CID135403798, CID344265, CID24858111, and CID155167581 generated a HOMO and LUMO energy score of -4.9072 and -1.9906, -5.14878 and -1.97464, -5.54058 and -2.38866, -5.33285 and -1.97164, and -5.4003 and -2.345 eV, respectively (Figure 2). Consequently, CID139030487 generated the lowest HOMO-LUMO gap (HLG) with the highest softness value of 0.69, where CID135403798, CID344265, CID24858111, and CID155167581 gave softness value of 0.65eV, 0.63eV, 0.63eV, 0.60eV, and 0.65eV respectively. We also calculated the chemical potential of these five compounds and found that these compounds have almost similar chemical potential ability. So, we selected these top five for pharmacophore modeling as they could be potential muti-target inhibitors.

**Figure 2.**
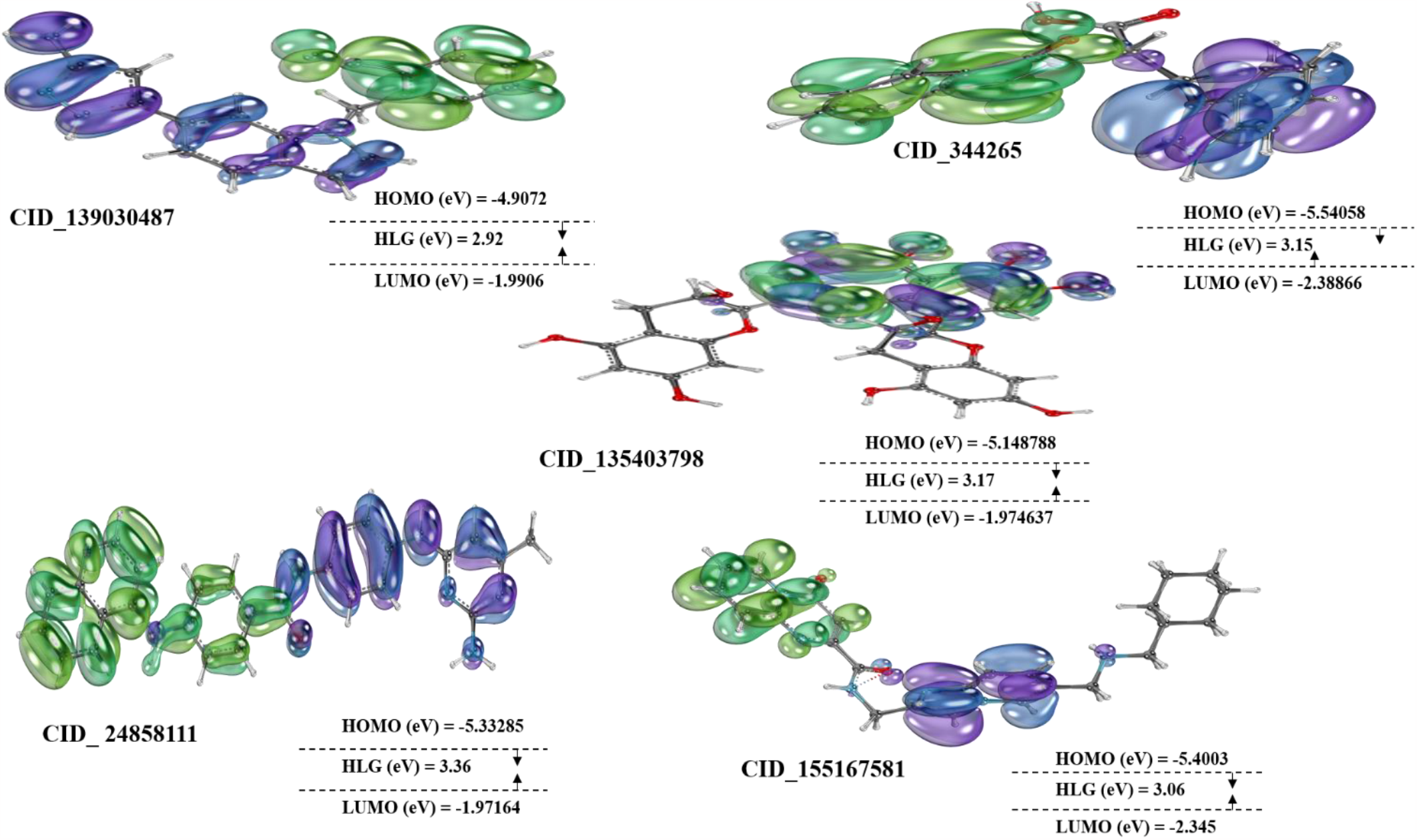
Frontier molecular orbital analysis of top 5 compounds. Here, HOMO (Highest occupied molecular orbitals) are in green while LUMO (Lowest unoccupied molecular orbitals) are in blue.

### 3.4 Pharmacophore modeling and Virtual screening of ZINC database

#### 3.4.1 Pharmacophore model generation and pharmacophore-based screening

At first, we generated a ligand structure-based pharmacophore model using top 5 selected multi-METTL targeting inhibitors with the highest frontier molecular orbital properties and then screened the ZINC database. Our pharmacophore model was generated by PharmaGIST, which predicted four spatial features (Aromatic-4, Hydrophobic-0, Donors-0, and Acceptors-0) in our generated model (S.figure 4). Then, the model was imported to ZINCPharmer to virtually screen the ZINC database. A maximum of 0.2 Å RMSD, 5 rotation bonds, and molecular weight between 350-550 from sphere centers were set as input parameters for ZINCPharmer, and a total of 483 hits was retrieved to make a ligand library for further molecular docking-based screening.

#### 3.4.2 Molecular Docking based screening

The hits of 483 compounds obtained in the pharmacophore-based screening were further screened against all five METTL proteins by AutoDock Vina tools to select the candidate compounds. After shortlisting the compounds according to the highest binding affinity score, binding pattern and interaction with individual METTL proteins were visualized by PyMOL to finalize the list of compounds that interact at the protein’s drug-binding catalytic active sites. Molecular docking and subsequent protein-ligand interaction site analysis demonstrated the 15 compounds with energetically top-scored (average binding energy > -10.0 kcal/mol) drug candidates for targeting METTL1, METTL3, METTL6, METTL16, and METTL18. Interestingly, further protein-ligand interaction analysis by PyMOL indicated that 13 out of above-mentioned 15 potent compounds have considerably occupied the catalytic SAM binding pockets of all METTL proteins. Among these 13 compounds, ZINC70666503 demonstrated the highest binding energy (13.1 kcal/mol) when interacting with METTL18. Intriguingly, the binding affinity of this compound to other METTL proteins was also notable (Table 2). On the other hand, ZINC40039977, ZINC25249143, and ZINC02659651 also exhibited notable binding affinity to all studied proteins. The docking results of all 483 compounds from the ZINC database are shown in S.table 5, and the top selected 13 compounds are shown in Table 2.

**Table 2:** Molecular docking results of top scored ZINC compounds with METTL family proteins

| **ZINC IDs** | **Molecular docking energy (kcal/mol)** |  |  |  |  | **Average** |
| --- | --- | --- | --- | --- | --- | --- |
|  | **METTL1** | **METTL3** | **METTL6** | **METTL16** | **METTL18** |  |
| ZINC09845394 | -8.9 | -10.2 | -10.5 | -10.2 | -11.1 | -10.18 |
| ZINC25249143 | -8.3 | -10.8 | -10.7 | -9.1 | -13 | -10.38 |
| ZINC27795763 | -8.6 | -9.8 | -11 | -8.9 | -12.7 | -10.2 |
| ZINC14891501 | -9.3 | -9.1 | -10.2 | -9.6 | -12.8 | -10.2 |
| ZINC40145195 | -9 | -8.6 | -10.8 | -10 | -12.8 | -10.24 |
| ZINC40039977 | -9.4 | -10.6 | -11 | -9.7 | -11.7 | -10.48 |
| ZINC33246308 | -9.2 | -9 | -11 | -10.3 | -11.8 | -10.26 |
| ZINC23632079 | -9.2 | -10.7 | -9.8 | -9.2 | -11.9 | -10.16 |
| ZINC70666503 | -9.4 | -10.4 | -11.1 | -9.1 | -13.1 | -10.62 |
| ZINC09124297 | -9.4 | -8.9 | -11.1 | -9.2 | -11.9 | -10.1 |
| ZINC13000658 | -9.5 | -9.8 | -10 | -9 | -11.8 | -10.02 |
| ZINC72324946 | -9.2 | -8.8 | -10.8 | -9.2 | -12.3 | -10.06 |
| ZINC02659651 | -9.6 | -11.9 | -11.7 | -11 | -9 | -10.64 |

NOTE: METTL1: Methyltransferase-like 1, METTL3: Methyltransferase-like 3, METTL6: Methyltransferase-like 6, METTL16: Methyltransferase-like 16, METTL18: Methyltransferase-like18

### 3.5 ADME and Toxicity prediction of top candidate inhibitors

The physicochemical properties, lipophilicity, water-solubility, and drug-likeness of 18 prospective drug candidates (CID139030487, CID135403798, CID344265, CID24858111, CID155167581 and 13 top-scored ZINC compounds) were evaluated by SwissADME (S.table 5). Among them, 3 compounds violated the Lipinski rule of five and showed undesired physiochemical properties in the case of molecular weight (MW), molecular refractivity (MR), and topological polar surface area (TPSA), which are considered as very important properties of a drug compound. Moreover, OSIRIS Property Explorer and admetSAR predicted the toxic and immunotoxic effects as well as cytochromes P450 (CYPs) isoforms inhibition of the several compounds. After initial ADMET properties screening, nine potential drug compounds with no predicted adverse effects were shortlisted (details are included in S.table 6). Finally, drug score prediction by OSIRIS Property Explorer predicted ZINC70666503 and ZINC13000658 as the top drug candidates, with drug scores of 87% and 82%. Additionally, ZINC70666503 has the blood barrier permeate ability with no AMES toxic effects (Table 3). Conversely, ZINC13000658 might be AMES toxic but not a blood-brain barrier permeable one which is important for drugs with a peripheral target as minimize BBB penetration ability might be required to reduce the possibility of undesired pharmacological events and avoid CNS side effects [65].

**Table 3:**
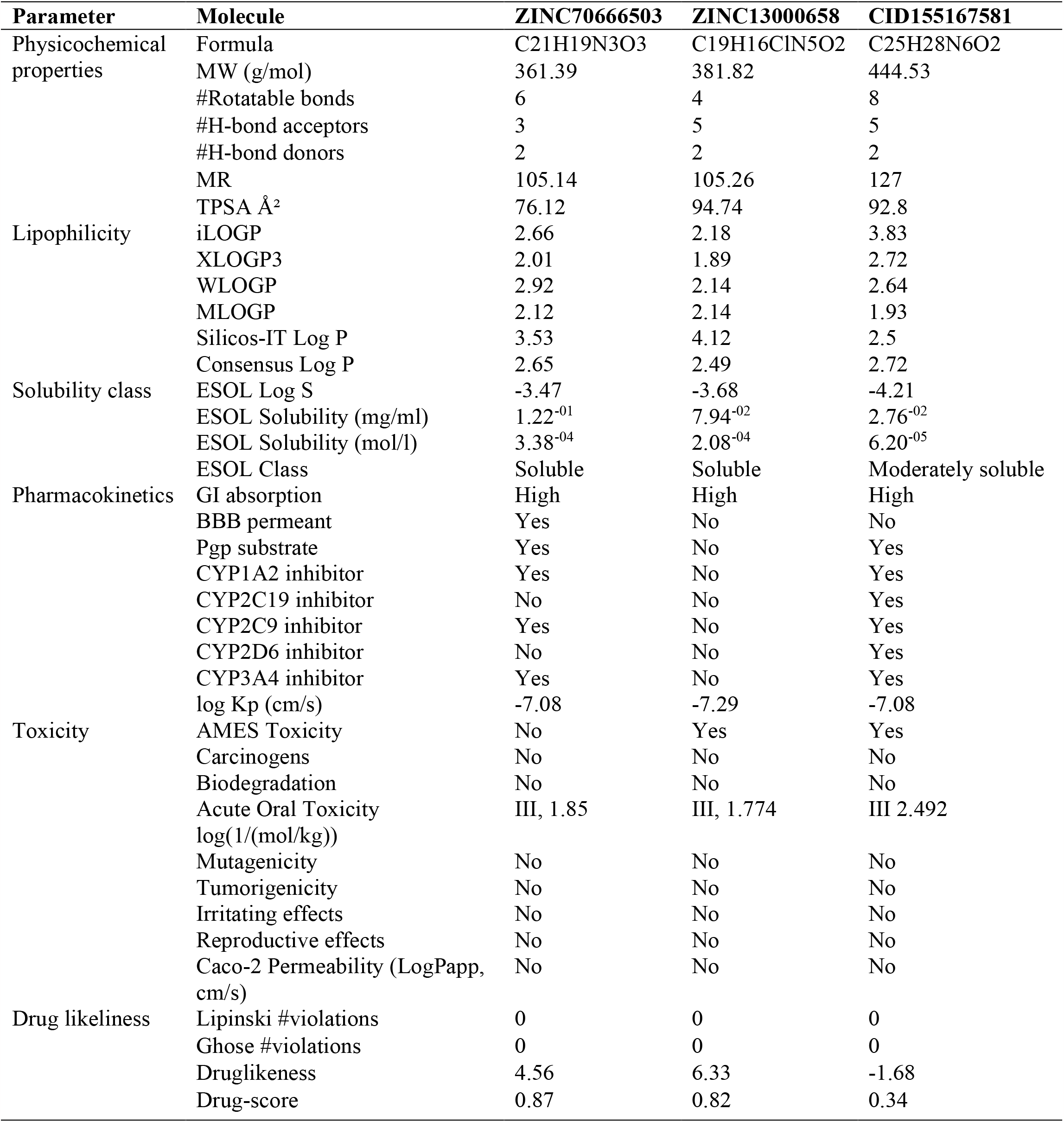
Absorption, Distribution, Metabolism, Excretion and Toxicity analysis of top 2 inhibitors

### 3.6 Re-docking and binding site analysis of candidate drugs

Site-specific protein-ligand re-docking of the final two drug compounds and CID155167581 against all METTL proteins was performed using AutoDock Vina followed by re-docking with two other docking server SwissDock, and CB Dock. Results indicated mostly similar results to the previous results with AutoDock Vina. In both AutoDock Vina and CB-Dock tools, the binding affinity score was found to be almost the same for all three compounds with previous docking results (S.table 7). In most cases, ZINC70666503 and ZINC13000658 had higher binding affinity than CID155167581 (STM2457), SAM competitive inhibitor of METTL3. Among our top 2 drug candidates, ZINC70666503 showed the lowest binding energy and the highest binding affinity to all METTL proteins. After docking, we identified the residues involved in the interaction and surprisingly found that both ZINC70666503 and ZINC13000658 interacted at the SAM binding catalytic domain of every METTL protein. As can be seen in Figure 3 and Figure 4, the interacted binding sites are at the interface of the studied METTL protein’s active site residues demonstrated earlier (Figure 1). Most surprisingly, binding site residues of the interacting sites of proteins-inhibitors complexes revealed that both ZINC70666503 and ZINC13000658 interacted with the amino acid residues that are the primary residues responsible for SAM binding at respective proteins’ active sites (Table 4).

**Figure 3.**
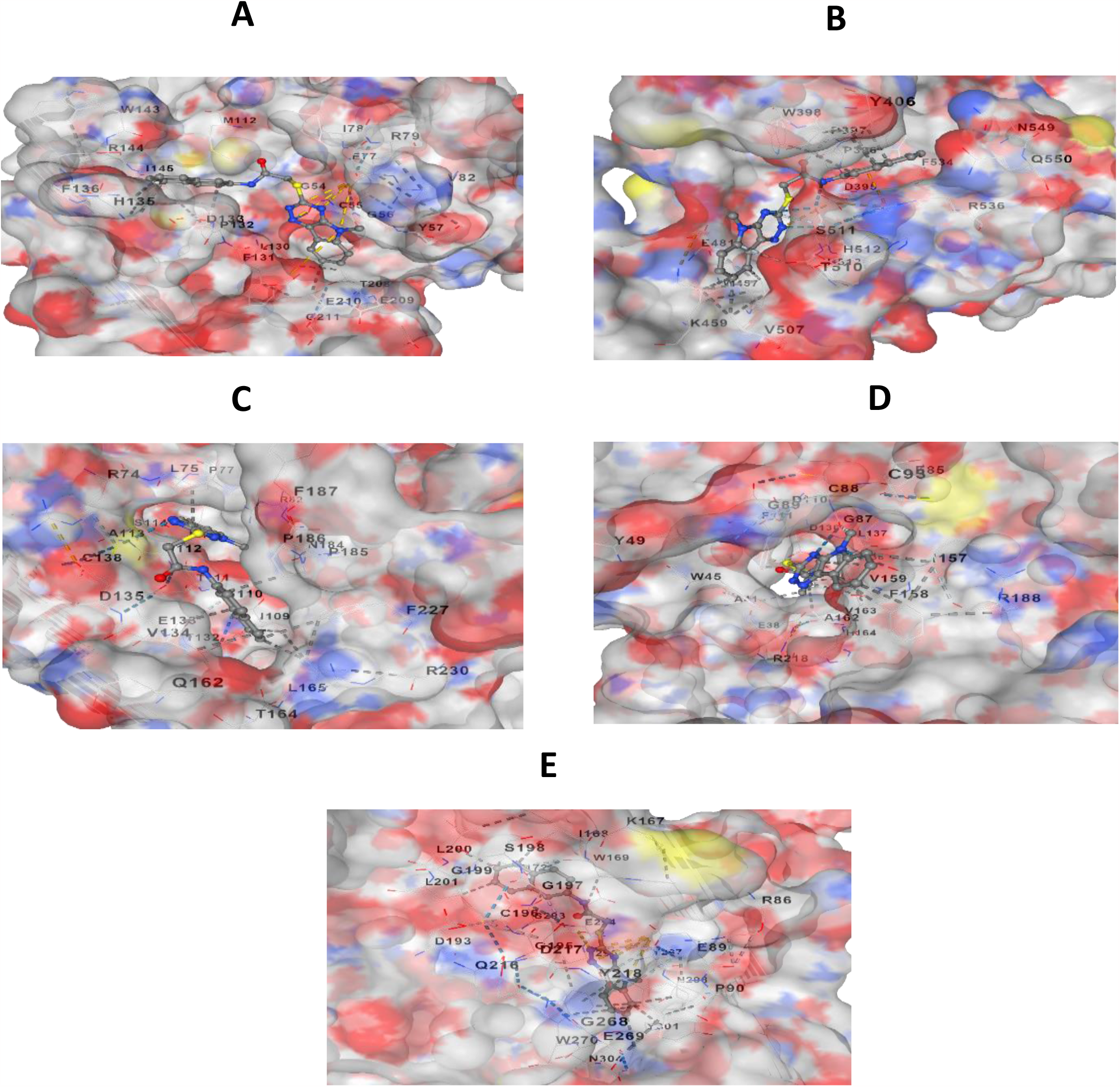
Binding site analysis of the interaction of ZINC70666503 at METTL1 (A), METTL3 (B), METTL6 (C), METTL16 (D), and METTL18 (E) proteins.

**Figure 4.**
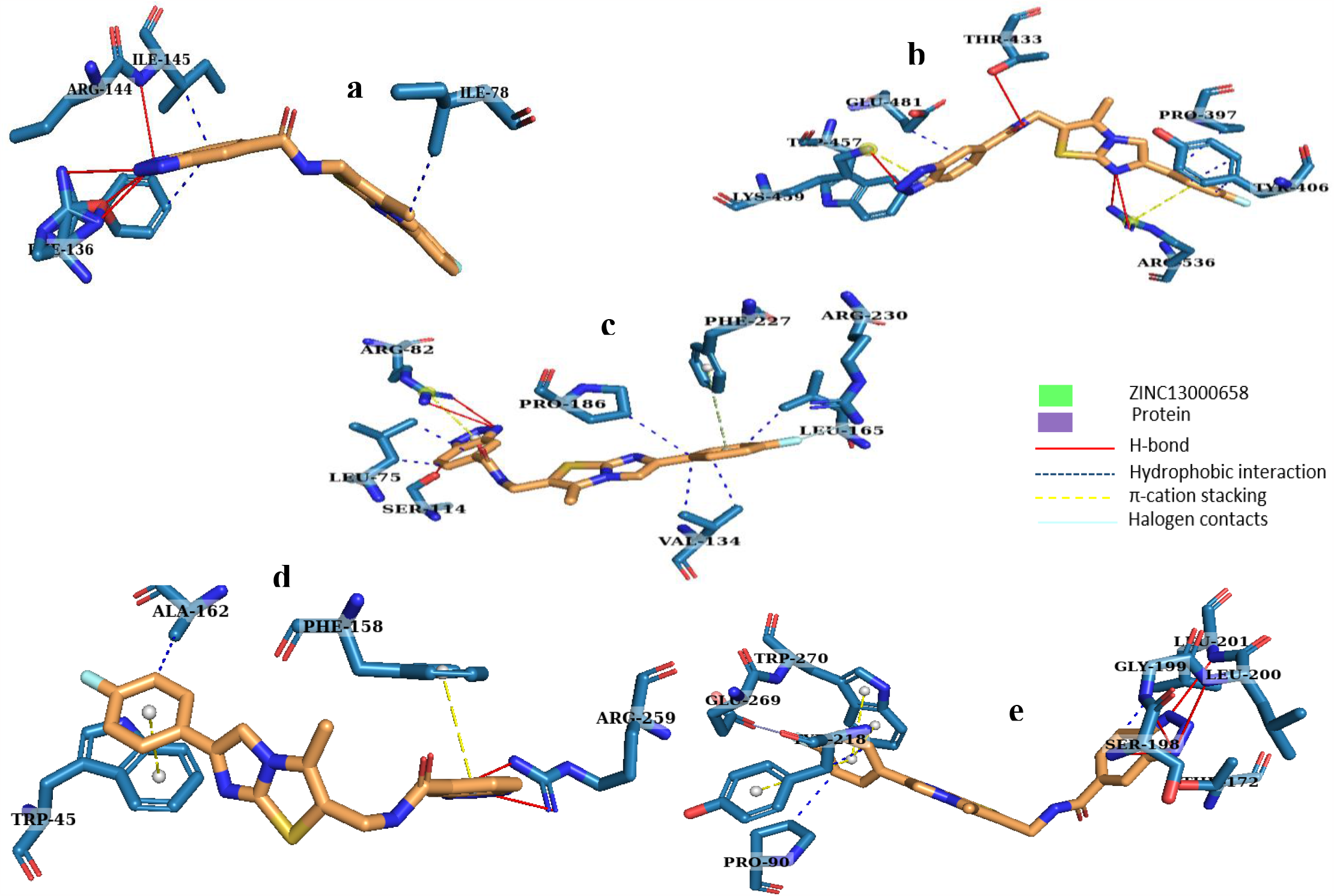
ZINC13000658 interacted with METTL1, METTL3, METTL6, METTL16, and METTL18 proteins by interacting with the amino acids shown in a, b, d, e, and c, respectively.

**Table 4.** Various interactions and interacting residues in the active pocket of methyltransferase family proteins with potential two drug candidates and S-adenosyl methionine

| Drug Candidates | METTL Proteins | H-bond | π-π stacking | Halogen bonds | Hydrophobic | Binding affinity (Kcal/mol) | Non-covalent bond |
| --- | --- | --- | --- | --- | --- | --- | --- |
| ZINC70666503 | METTL1 | - | - | - | 57Y, 129F, 133D, 145I, 209E, 536R | -9.4 | 6 |
|  | METTL3 | 511S, 536R | 534F | - | 378I, 397P, 409L, 433T, 534F, 549N | -10.4 | 12 |
|  | METTL6 | 159V | 111F | - | 41A, 138T, 157I, 158F, 159V, 162A, 163V | -9.4 | 11 |
|  | METTL16 | 75L | 227F | - | 75L, 109I, 134V, 165L, 186P | -9.1 | 8 |
|  | METTL18 | 197G, 297Y | 270W | - | 90P, 168I, 172T, 200L, 201L, 218Y, 269E | -13.1 | 13 |
| ZINC13000658 | METTL1 | 136F, 144R, 145I | - | - | 78I, 136F, 144R, 145I | -9.5 | 7 |
|  | METTL3 | 433T, 459K, 536R | 459K, 536R | - | 397P, 406Y, 457W, 481E | -9.8 | 11 |
|  | METTL6 | 259R | 45W, 158F | - | 162A | -10 | 5 |
|  | METTL16 | 82R, 114S | 227F | 230R | 75L, 134V, 165L, 186P | -9 | 12 |
|  | METTL18 | 172T, 198S, 199G, 200L, 201L | 218Y, 270W | 269E | 90P, 201L, 218Y | -11.8 | 12 |
| S-Adenosyl Methionine | METTL1 | 54G, 55C, 77G, 78I, 133D, 143W, 145I, 209E, 210E | - | - | 78I | -7.5 | 9 |
|  | METTL3 | 377D, 378I, 395D, 536R, 549N, 550Q | - | - | 513K | -7.7 | 10 |
|  | METTL6 | 49Y, 85E, 90V, 92N, 159V, 190Y, 213R | - | - | 45W, 190Y | -7.7 | 13 |
|  | METTL16 | 75L, 82R, 110G, 114S, 134V | - | - | 134V | -7.3 | 9 |
|  | METTL18 | 169W, 172T, 195G, 196C, 197G, 198S, 216Q, 218Y, 269E, 270W | 218Y, 270W | - | 89E | -10.5 | 16 |

NOTE: METTL1: Methyltransferase-like 1, METTL3: Methyltransferase-like 3, METTL6: Methyltransferase-like 6, METTL16: Methyltransferase-like 16, METTL18: Methyltransferase-like18; A: Alanine, R: Arginine, N: Asparagine, D: Aspartic acid, C: Cysteine, E: Glutamic acid, Q: Glutamine, G: Glycine, H: Histidine, I: Isoleucine, L: Leucine, K: Lysine, M: Methionine, F: Phenylalanine, P: Proline, S: Serine, T: Threonine, W: Tryptophan, Y: Tyrosine, V: Valine; SAM: S-Adenosyl Methionine

### 3.7 Molecular Dynamics Simulation

Although molecular docking is a widely-used techniques, reliable predictions for binding affinities, drug-receptor intermolecular interactions, solvent effects, dynamics must be considered. Therefore, molecular dynamics simulation for the METTL1, METTL3, METTL6, METTL16, and METTL18 in complex with ZINC70666503 and ZINC13000658 was performed to determine the stability and rigidity of protein-ligand complexes in a specific artificial cellular environment at nanosecond scaled. Root mean square deviation (RMSD) of METTL proteins and ZINC70666503 showed a constant binding pattern and no significant fluctuation during the 100ns time frame. However, a slight fluctuation was observed for RMSD of METTL6 near 70 ns when complexed with ZINC13000658. However, primarily stable conditions with no dramatic inconstancy had been observed in the RMSD of all METTL proteins in complex with these two inhibitors (Figure 5). Besides, flexibility estimation of protein and ligand complex by RMSF indicated that the fluctuations were confined in the 1 nm range, which indicates higher stability. On the other hand, the lower Rg value indicates high compactness, and the larger value evidences the dissociation of the inhibitors from the protein. For both ZINC70666503 and ZINC13000658, lower Rg values were also measured. Moreover, H-bond estimation during this simulation time showed ZINC13000658 had a maximum number of 7 hydrogen bonds when interacting with METTL18. This ligand also exhibited a more significant number of interacting H-bond numbers than ZINC70666503 with other METTL proteins. An average of 4 hydrogen bonds formed during METTL protein’s interactions with ZINC13000658 and two hydrogen bonds between METTL proteins and ZINC70666503 interactions (S.figure 5).

**Figure 5.**
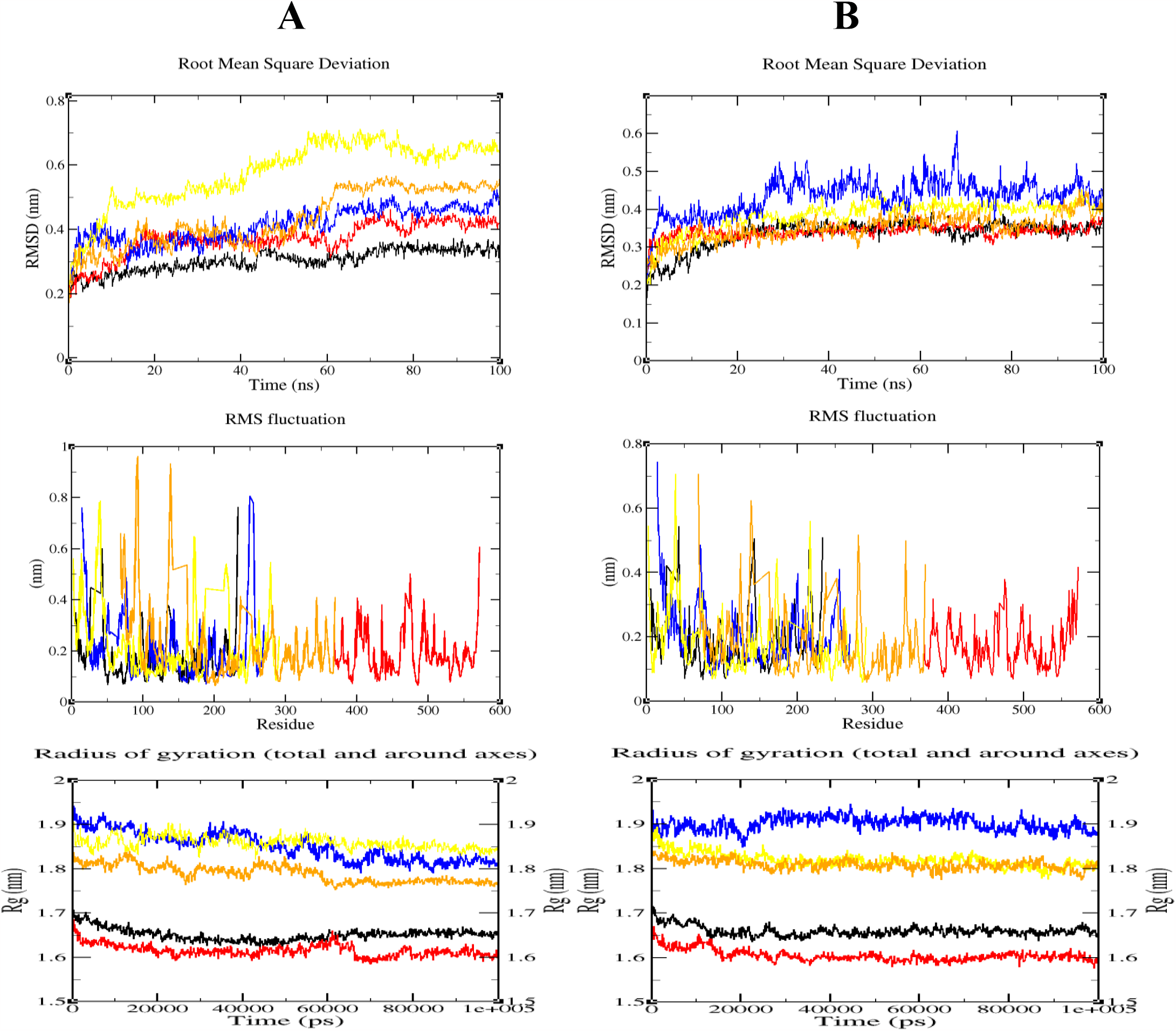
Root Mean Square Deviation (RMSD), Root Mean Square Fluctuation (RMSF), and radius of gyration (Rg) of all studied METTL proteins in complex with ZINC70666503 (left, A) and ZINC13000658 (right, B). Here, black, red, blue, yellow, and orange color denote for METTL1, METTL3, METTL6, METTL16, and METTL18, respectively.

### 3.8 MM/PB(GB)SA analysis from post molecular dynamics trajectory

We adopted MM/PB(GB)SA analysis to calculate the ligand-binding free energy to the desired protein by calculating electrostatic energy (ELE), Van der Waals contribution (VDW), and total binding energy. This energy calculation showed that the value of total binding energy in the METTL18 in complex with ZINC70666503 is -355.124 kcal/mol, which is higher than the other complexes for this compound (Table 5). On the other hand, ZINC13000658 had a value of -388.081 kcal/mol when complexed with METTL6. Most importantly, except for that of the METTL1-ZINC70666503 complex, most of the protein-ligand complex arguably showed significant total binding energy (>-300kcal/mol) for over the 100 ns simulation time (Table 5).

**Table 5.** MM-PB(GB)SA binding free energy analysis

| Drug Candidates |  | Energy value (kcal/mol) |  |  |  |  |
| --- | --- | --- | --- | --- | --- | --- |
|  |  | METTL1 | METTL3 | METTL6 | METTL16 | METTL18 |
| ZINC70666503 | ESE | 3.705 | -41.771 | -24.415 | -33.540 | -33.416 |
|  | VDW | -229.629 | -280.344 | -318.267 | -306.897 | -321.708 |
|  | Total binding energy | -225.924 | -322.115 | -342.682 | -340.437 | -355.124 |
| ZINC13000658 | ESE | -67.249 | -52.096 | -90.336 | -55.118 | -59.652 |
|  | VDW | -264.633 | -256.243 | -297.745 | -240.819 | -310.378 |
|  | Total binding energy | -331.882 | -308.339 | -388.081 | -295.937 | -370.030 |

NOTE: ESE: Electrostatic energy, VDW: Van der waals energy, METTL1: Methyltransferase-like 1, METTL3: Methyltransferase-like 3, METTL6: Methyltransferase-like 6, METTL16: Methyltransferase-like 16, METTL18: Methyltransferase-like18;

## 4. DISCUSSION

Development and progression of HCC is a complex process. Inactivation of multiple tumor suppressor gene, abnormal activation of oncogenes (K-ras, BRAF, etc.), frequent alteration of tyrosine kinase and growth factor receptors, and different signalling pathways are the key players in this complex process [66]. Therefore, different targeted and immunotherapeutic treatment strategies are employed targeting biomarkers involved in those HCC promoting key signaling pathways [67]. However, poor treatment efficiency and drug resistance phenomenon of different targeted and immunotherapeutic treatment urge the development of efficient strategies and potential therapeutic agents along with druggable targets to increase the pharmacological treatment sensitivity of HCC [8,9].

Recent studies have demonstrated the central role of methyltransferase like proteins METTL1, METTL3, METTL6, METTL16, and METTL18 in regulating aberrant expression in human hepatocellular carcinoma tissues and pathogenesis and progression of HCC cells, such as cell survival, proliferation, metastasis, and invasion which in turns makes them novel biomarkers [11-17]. However, despite identifying a few multi-kinases or receptors targeting anti-cancer inhibitors, no multi-target drugs or inhibitors have been reported against HCC-associated METTL proteins [68]. Therefore, we performed to discover effective and highly specific first-in-class multi-targeting Methyltransferase-like proteins inhibitors by repurposing previously identified SAM competitive inhibitors or inhibitors targeting the active site of the SAM binding pockets of microbial tRNA methyltransferase, human DNMT and METTL3 proteins. We employed comprehensive molecular docking-based screening, extensive structural and protein-ligand interaction analysis, DFT analysis, pharmacophore modeling, and molecular dynamics simulation of the potent small molecule inhibitors.

Molecular docking and studying drug-binding active site have been substantially used in the drug discovery process to predict binding site complementarity between a therapeutic target and a drug ligand and to understand the molecular interaction between the target and drug candidates [69, 70]. Therefore, towards discovering potent hepatocellular carcinoma-associated METTL proteins inhibitors, 73 selected small molecule inhibitors were screened through molecular docking and comprehensive binding site analysis of top five compounds in complex with respective METTL proteins with the highest binding energy indicated similar binding patterns that might indicate drug-binding active sites of respective METTL proteins. For instance, high docking scores for top ligands with individual METTL proteins are attributed to their ability to form a complex at the respective METTL protein’s S-adenosyl-L-methionine (SAM) binding cavity (Figure 1). Moreover, our METTL3-ligands interaction analysis (Table 1) supports the findings that have been reported in the literature of Yankova et al. 2021, where they illustrated amino acids residues with positions at 378I, 395D, 406Y, 431W, 457W, 511K, 513K, 532E, 534F, 536R, and 549N as the key active site residues for forming METTL3-STM2457 complex [18]. Other than that, for the first time, the present study identifies key amino acid residues of METTL1, METTL6, METTL16, and METTL18 proteins’ catalytic sites involved in interaction with potent drug compounds (Table 1).

As we focused on inhibiting more than one protein, compounds with higher chemical reactivity and softness value would be highly capable of interacting with multiple target proteins [71]. So, we adopted frontier molecular orbital properties analysis which is one of the essential methods for determining the pharmacological properties of drug compounds (intramolecular and intermolecular charge transfer, HOMO/LUMO interactions, chemical hardness, softness, and chemical potentials) [48, 72]. After calculating the softness and chemical potentiality from EHOMO-ELUMO values, we had five compounds (2 tRNA methyltransferase inhibitors, 2 DNMT inhibitors, and 1 METTL3 inhibitor) with softness values of greater than 0.6 eV (S.table 4), suggesting the compounds’ considerably higher chemical instability and high chemical reactivity in contrast to others. Then, we generated a ligand-based pharmacophore model (S.figure 4) and screened the ZINC database using the model to identify chemical compounds with comparable pharmacophore features as this has potentiality to screen large libraries for identifying hits with specific desired features [73]. Subsequently, we further docked the hits of 483 compounds against all five METTL proteins and conducted protein-ligand interaction analysis to identify the compounds with higher binding affinity at the proteins’ SAM binding catalytic sites.

Finally, we adopted ADMET analysis-based filtration approaches to finalizing the selection. Computational ADME/T properties analysis of the top two drug candidates, ZINC70666503 and ZINC13000658, showed minimum cytochromes P450 (CYP) isoforms inhibition, highest drug scores, and no predicted toxic effects in comparison to STM2457 (Table 5). Cytochrome P450s enzymes, particularly CYP3A4, CYP2D6, CYP2C9, and CYP2C19, are important for drug metabolism in humans; for instance, CYP3A4 inactivation may enhance drug toxicity through increased exposure to other co-administered drugs [74,75]. In that case, ZINC13000658 has no CYP isoform inhibition activity, whereas ZINC70666503 may interact with one or two isoforms. To ensure the reliability of molecular docking, we re-evaluated the docking performance of our top 2 drug candidates using three docking tools, and significant concordance was observed between site-specific docking by AutoDock Vina and re-docking results of CB-Dock with the previous AutoDock Vina results. Notably, both ZINC70666503 and ZINC13000658 interacted with the amino acid residues of the pocket of the adenosine moiety of SAM (Table 4), which might be the reason for higher interaction with SAM-dependent methyltransferase [18,36].

Our present study is the first to identify ZINC70666503 and ZINC13000658 (S.figure 2) as METTL1, METTL6, METTL16, and METTL18 inhibitors. Although some potent METTL3 inhibitors such as STM2457, Quercetin, and Eltrombopag have been studied recently, and their anti-proliferative effects, increase apoptosis and reduced growth in the acute myeloid leukemia (AML) cell line have also been demonstrated, their adverse reaction and drug safety on treating AML remain unknown [35]. Our computational study on molecular docking and ADMET analysis not only predicted higher binding affinity of ZINC70666503 and ZINC13000658 but also indicated that they have higher pharmacokinetic, drug likeliness properties with no undesirable toxic effects compared to STM2457, Quercetin, UZH1a, Eltrombopag and other METTL3 inhibitors (S.table 3 and Table 3).

We also assessed the stability of both five METTL proteins-ZINC70666503 complexes and proteins-ZINC13000658 complexes by measuring ligand-receptor RMSD, RMSF, radius of gyration, number of H-bonds, and calculation of MM/PB(GB)SA binding free energies because trajectories obtained from molecular dynamics simulation could be used to predict the stability of the protein-inhibitors complexes in an aqueous solution by analyzing the best-docked pose predicted through the docking analysis [76]. The results indicated the stability of most of the protein-inhibitor complexes over the entire simulation time and the inhibitory potential of both inhibitors, which are due to efficient binding to the catalytic drug binding sites of METTL1, METTL3, METTL6, METTL16, and METTL18.

Moreover, the study also points out the higher selectivity of these two compounds towards all five SAM-dependent HCC-associated METTL proteins in a SAM-competitive inhibition mechanism. Notably, residues involved in hydrogen bonding (H-bond) of two inhibitors-proteins complexes indicate that ZINC13000658 formed higher number of H-bond compared to ZINC70666503 (Table 4) and might hamper the hydrogen bond formation between SAM and METTL. Overall, these findings provided evidence for the two identified SAM-competitive catalytic inhibitors of METTL proteins as promising anti-cancer drug candidates for hepatocellular carcinoma, which need further investigations of their specificity to METTL proteins. Moreover, to overcome cancer drug resistance and provide new treatments, our predicted two compounds can also be combined with the existing first-line and second-line therapies for hepatocellular carcinoma as this can enhance the efficacy of therapies mediated by targeted drugs [77].

Additionally, our studied five METTL proteins are associated with not only hepatocellular carcinoma but also different human cancers. For example, methyltransferase-like 1 (METTL1) is overexpressed in lung cancer, bladder cancer, lung adenocarcinoma, head and neck squamous cell carcinoma, and gastric cancer [78-82]. On the other hand, methyltransferase-like 3 (METTL3) is upregulated in lung adenocarcinoma, breast carcinoma, triple-negative breast cancer, and several other cancers. [83]. In that case, ZINC70666503 (EiM08-22770) and ZINC13000658 (AKOS033010880) could also be promising therapeutics for other deadly human cancers.

## 5. Conclusion

Hepatocellular carcinoma is one the most prominent health issue worldwide with no effective treatment strategies. In this study, two compounds ZINC70666503 and ZINC13000658 showed promising results that could expedite multi-targeting drug development against five METTL proteins involved in HCC. ZINC70666503 and ZINC13000658 could be more promising options for treating human malignancies than any previously identified METTL3 inhibitors. Remarkably, these two compounds are the first identified catalytic inhibitors that can act against METTL1, METTL6, METTL16, and METTL18. However, further *in vivo* efficacy testing and exploration of their anti-tumor activity are highly needed.

## Key Points

- ZINC70666503 and ZINC13000658 were identified as the novel potential multi-target inhibitors of methyltransferase like protein METTL1, METTL3, METTL6, METTL16, and METTL18 which play roles in hepatocellular carcinoma progression through pharmacophore modeling and ZINC database screening.
- These two compounds (with 87% and 82% drug scores) interacted with the amino acid residues that are responsible for S-Adenosyl Methionine (SAM) binding at the active site cavity of proteins and might be SAM competitive inhibitors.
- RMSD, RMSF, radius of gyration, and H-bond analysis supported the binding stability of protein-inhibitor complexes over the 100ns simulation time.

## Ethical approval

Not required.

## Data availability

All data supporting the findings of this study are available within the article and its supplementary materials.

## Author contributions

**Md. Niaz Morshed:** Conceptualization, Methodology, Software, Formal analysis, Investigation, and Data interpretation, Writing - original draft, and review & editing. **Md. Sorwer Alam Parvez:** Methodology, Data interpretation, Writing - review & editing. **Rakibul Islam Akanda:** Software, Formal analysis, Writing - review & editing. **Manash Kumar Saha:** Software, Formal analysis, Writing - review & editing. **Jannatul Fardous:** Formal analysis, Writing - review & editing. **Mohammad Jakir Hosen:** Project administration, Supervision, Writing - review & editing.

## Author Information

**Md. Niaz Morshed** is a Research Assistant at Department of Genetic Engineering and Biotechnology, Shahjalal University of Science and Technology. He is interested in molecular mechanisms of cancer progression, and therapeutic resistance and drug development.

**Md. Sorwer Alam Parvez** is a Research Student at Kyoto University Graduate School in Medicine.

**Rakibul Islam Akondo** is an undergraduate student, majoring in Genetic Engineering and Biotechnology at the Shahjalal University of Science and Technology. His research interest is bioinformatics, cancer biology.

**Manash Kumar Saha** is currently majoring in Genetic Engineering and Biotechnology at Shahjalal University of Science and Technology. He is mainly interested in immunoinformatics, virology, vaccinology and evolutionary biology.

**Jannatul Fardous** has completed her Bachelor of Medicine and Bachelor of Surgery (MBBS) from Chittagong Medical College, Chittagong, Bangladesh. Her research focus is inclined to medical oncology and immunology.

**Mohammad Jakir Hosen** is a Professor and PI of the Genetic Disease and Public Health research group of the Department of Genetic Engineering and Biotechnology, Shahjalal University of Science and Technology, Sylhet-3114, Bangladesh.

