## Supplementary material for "Small Molecule Inhibitors Targeting Methyltransferase-Like (METTL) Proteins Against Hepatocellular Carcinoma: A Comprehensive Drug Repurposing Approach": S.table 1

**S.table 1. Selected drug candidate list**

| No. | Pubchem CID No. | Chemical Name |
|---|---|---|
| CID1 | CID135403798 | Theaflavin |
| CID2 | CID137347756 | N-[(4-{[cyclohexyl(ethyl)amino]methyl}phenyl)methyl]-4-oxo-3,4-dihydrothieno[2,3-d]pyrimidine-5-carboxamide |
| CID3 | CID97429072 | N-[(3S)-1-(5-pyridin-4-yl-1H-pyrazol-3-yl)piperidin-3-yl]-1H-indole-2-carboxamide |
| CID4 | CID137347756 | N-[(4-{[cyclohexyl(ethyl)amino]methyl}phenyl)methyl]-4-oxo-3,4-dihydrothieno[2,3-d]pyrimidine-5-carboxamide |
| CID5 | CID137347757 | N-({4-[(octylamino)methyl]phenyl}methyl)-4-oxo-3,4-dihydrothieno[2,3-d]pyrimidine-5-carboxamide |
| CID6 | CID137347758 | N-({4-[(diethylamino)methyl]phenyl}methyl)-4-oxo-3,4-dihydrothieno[2,3-d]pyrimidine-5-carboxamide |
| CID7 | CID71724899 | N-[[4-[(4-aminopiperidin-1-yl)methyl]phenyl]methyl]-4-oxo-3H-thieno[2,3-d]pyrimidine-5-carboxamide |
| CID8 | CID139030490 | 5-azanyl-3-[1-(pyridin-4-ylmethyl)indol-6-yl]-1~{H}-pyrazole-4-carbonitrile |
| CID9 | CID2734361 | Indole-5-boronic acid |
| CID10 | CID139030485 | 3-[1-[(4-methoxyphenyl)methyl]indol-6-yl]-1~{H}-pyrazol-5-amine |
| CID11 | CID 139030486 | 5-azanyl-3-(1~{H}-indol-6-yl)-1~{H}-pyrazole-4-carbonitrile |
| CID12 | CID139030487 | 2-[[6-(5-azanyl-1~{H}-pyrazol-3-yl)indol-1-yl]methyl]benzenecarbonitrile |
| CID13 | CID139030488 | 3-[1-(phenylmethyl)indol-6-yl]-1~{H}-pyrazol-5-amine |
| CID14 | CID139030489 | 5-azanyl-3-[1-(pyridin-3-ylmethyl)indol-6-yl]-1~{H}-pyrazole-4-carbonitrile |
| CID15 | CID139030490 | 5-azanyl-3-[1-(pyridin-4-ylmethyl)indol-6-yl]-1~{H}-pyrazole-4-carbonitrile |
| CID16 | CID139030491 | 3-azanyl-5-[3-chloranyl-1-(pyridin-3-ylmethyl)indol-6-yl]-1~{H}-pyrazole-4-carbonitrile |
| CID17 | CID139030492 | 5-azanyl-3-[1-[(2-oxidanylpyridin-3-yl)methyl]indol-6-yl]-1~{H}-pyrazole-4-carbonitrile |
| CID18 | CID139030493 | 5-azanyl-3-[1-[[(3~{S})-1-methylpiperidin-3-yl]methyl]indol-6-yl]-1~{H}-pyrazole-4-carbonitrile |
| CID19 | CID145946018 | -amino-5-[1-[[(2R)-1-methylpiperidin-2-yl]methyl]indol-6-yl]-1H-pyrazole-4-carbonitrile |
| CID20 | CID139030495 | 5-azanyl-3-[1-[[4-(piperidin-1-ylmethyl)phenyl]methyl]indol-6-yl]-1~{H}-pyrazole-4-carbonitrile |
| CID21 | CID139030496 | 5-azanyl-3-[1-[[4-(morpholin-4-ylmethyl)phenyl]methyl]indol-6-yl]-1~{H}-pyrazole-4-carbonitrile |
| CID22 | CID139030497 | 3-[1-[[4-(piperidin-1-ylmethyl)phenyl]methyl]indol-6-yl]-1~{H}-pyrazol-5-amine |
| CID23 | CID145997887 | 5-azanyl-3-[1-[[(3~{R})-1-(phenylmethyl)piperidin-3-yl]methyl]indol-6-yl]-1~{H}-pyrazole-4-carbonitrile |

| | | |
|---|---|---|
| CID24 | CID139030498 | 3-[1-[(3-methoxyphenyl)methyl]indol-6-yl]-1~{H}-pyrazol-5-amine |
| CID25 | CID139030500 | 5-azanyl-3-[1-[[4-[(4-propan-2-ylpiperazin-1-yl)methyl]phenyl]methyl]indol-6-yl]-1~{H}-pyrazole-4-carbonitrile |
| CID26 | CID139030501 | (3-methoxyphenyl)methyl 1~{H}-pyrazole-4-carboxylate |
| CID27 | CID68910199 | Benzyl 1H-pyrazole-4-carboxylate |
| CID28 | CID139030502 | [3-(2-methoxy-2-oxidanylidene-ethoxy)phenyl]methyl 1~{H}-pyrazole-4-carboxylate |
| CID29 | CID71677803 | N-(4-{[(1h-Imidazol-2-Ylmethyl)amino]methyl}benzyl)-4-Oxo-3,4-Dihydrothieno[2,3-D]pyrimidine-5-Carboxamide |
| CID30 | CID71677804 | N-[4-(Aminomethyl)benzyl]-4-Oxo-3,4-Dihydrothieno[2,3-D]pyrimidine-5-Carboxamide |
| CID31 | CID215639 | 5-Phenyl-3H-thieno[2,3-d]pyrimidin-4-one |
| CID32 | CID56703871 | (2-hydroxy-N-({1-[2-hydroxy-1-(hydroxymethyl)ethyl]piperidin-3-yl}methyl)-5-methylbenzamide) |
| CID33 | CID56900216 | (N-(2-isobutoxybenzyl)-N,2-dimethyl-2,8-diazaspiro[4.5]decane-3-carboxamide) |
| CID34 | CID23306 | Coralyne, |
| CID35 | CID4501 | Nitidine |
| CID36 | CID3025944 | Lomeguatrib |
| CID37 | CID65482 | Sinefungin |
| CID38 | CID5567 | Trifluperidol |
| CID39 | CID5494506 | Inauhzin |
| CID40 | CID702558 | N-Phthaloyl-L-tryptophan (RG108) |
| CID41 | CID9444 | 5-azacytidine |
| CID42 | CID451668 | Decitabine |
| CID43 | CID100016 | Zebularine |
| CID44 | CID4914 | Procaine |
| CID45 | CID65064 | (K)-epigallocatechin-3-gallate (EGCG) |
| CID46 | CID6400741 | Pasmmaplin A |
| CID47 | CID344265 | NSC401077 (RG108) |
| CID48 | CID327606 | NSC303530 |
| CID49 | CID5270713 | 5-Aza-4'-thio-2'-deoxycytidine |
| CID50 | CID132233544 | (2R)-2-{[6-(4-aminopiperidin-1-yl)-3,5-dicyano-4-ethylpyridin-2-yl]sulfanyl}-2-phenylacetamide |
| CID51 | CID3637 | Hydralazine |

| | | |
|---|---|---|
| CID52 | CID25150857 | BIX-01294 |
| CID53 | CID442757 | Nanaomycin A |
| CID54 | CID47751 | Fazarabine |
| CID55 | CID135564655 | Guadecitabine |
| CID56 | CID10037425 | 4′-thio-2′-deoxycytidine (tdcyd) |
| CID57 | CID515328 | 5-Fluoro-2′-deoxycytidine (fdcyd) |
| CID58 | CID24858111 | SGI-1027 |
| CID59 | CID76318201 | MC3343 |
| CID60 | CID156587340 | 6-[4-[6-[(4,4-dimethylpiperidin-1-yl)methyl]pyridin-3-yl]-1-oxa-4,9-diazaspiro[5.5]undecan-9-yl]-N-(phenylmethyl)pyrimidin-4-amine |
| CID61 | CID11271909 | 4-[2-[5-chloro-1-(diphenylmethyl)-2-methyl-1H-indol-3-yl]-ethoxy]benzoic acid (CDIBA) |
| CID62 | CID135449332 | Eltrombopag |
| CID63 | CID5280343 | Quercetin |
| CID64 | CID155167581 | STM2457 |
| CID65 | CID154815692 | 4-[(4,4-dimethylpiperidin-1-yl)methyl]-2-oxidanyl-~{N}-[[(3~{R})-3-oxidanyl-1-[6-[(phenylmethyl)amino]pyrimidin-4-yl]piperidin-3-yl]methyl]benzamide |
| CID66 | CID60961 | Adenine riboside |
| CID67 | CID98567 | 5'-N-methylcarboxamidoadenosine |
| CID68 | CID97692 | Adenosine 5'-carboxamide |
| CID69 | CID110145076 | (2~{S},3~{S},4~{R},5~{R})-5-(6-aminopurin-9-yl)-3,4-bis(oxidanyl)-~{N}-piperidin-4-yl-oxolane-2-carboxamide |
| CID70 | CID110154059 | HMS3847E10 |
| CID71 | CID145994381 | (2~{R},3~{R},4~{R},5~{R})-2-[(6-aminopurin-9-yl)methyl]-5-azanyl-oxane-3,4-diol |
| CID72 | CID440041 | Cpd-564 |
| CID73 | CID156587337 | 4-[(4,4-dimethylpiperidin-1-yl)methyl]-~{N}-[[(3~{R})-1-[6-(methylamino)pyrimidin-4-yl]-3-oxidanyl-piperidin-3-yl]methyl]benzamide |

### S.table 2. Grid box size and center for molecular docking study

| Grid Box | METTL1 | METTL3 | METTL6 | METTL16 | METTL18 |
|---|---|---|---|---|---|
| size (x, y, z) | 98Å × 96Å × 94Å | 56 Å × 72 Å × 68 Å | 64 Å × 56 Å × 58 Å | 64 Å × 56 Å × 38 Å | 30 Å × 60 Å × 44 Å |
| center (x, y, z) | 38.808 Å × -11.89 Å × -4.947 Å | -43 Å × 31 Å × 26 Å | 41.481 Å × -19.234 Å × 12.272 Å | -9.956 Å × 9.819 Å × 27.567 Å | 8.338 Å × 4.937 Å × 29.508 Å |

NOTE: METTL1: Methyltransferase-like 1, METTL3: Methyltransferase-like 3, METTL6: Methyltransferase-like 6, METTL16: Methyltransferase-like 16, METTL18: Methyltransferase-like18

### S.table 3. Molecular docking results of selected 73 compounds

| | Binding energy (Kcal/mol) | | | | | |
|---|---|---|---|---|---|---|
| **Pubchem CID No.** | **METTL1** | **METTL3-14** | **METTL6** | **METTL16** | **METTL18** | **Average** |
| CID135403798 | -10.3 | -9.2 | -9.1 | -10.2 | -8.2 | -9.4 |
| CID137347756 | -8.1 | -8 | -8.2 | -8.3 | -11.1 | -8.74 |
| CID97429072 | -8.5 | -8.2 | -10.7 | -9.2 | -7.3 | -8.78 |
| CID137347756 | -8.6 | -7.9 | -7.8 | -8.3 | -7 | -7.92 |
| CID137347757 | -6.7 | -7 | -7.4 | -7.5 | -5.5 | -6.82 |
| CID137347758 | -7.5 | -7.4 | -7.1 | -7.6 | -6.3 | -7.18 |
| CID71724899 | -8.2 | -8 | -7.5 | -7.6 | -11.1 | -8.48 |
| CID139030490 | -7.6 | -8.4 | -7.7 | -8 | -7.1 | -7.76 |
| CID2734361 | | | | | | 0 |
| CID139030485 | -7.8 | -8.9 | -9 | -8.2 | -7.5 | -8.28 |
| CID 139030486 | -6.9 | -7.4 | -7.1 | -7.2 | -9.6 | -7.64 |
| CID139030487 | -8.6 | -8.6 | -8.7 | -8.3 | -10.8 | -9 |
| CID139030488 | -8.7 | -9 | -11.3 | -8.7 | -7.7 | -9.08 |
| CID139030489 | -8.2 | -7.7 | -7.7 | -8 | -7.4 | -7.8 |
| CID139030490 | -8 | -8.4 | -7.7 | -8 | -10.3 | -8.48 |
| CID139030491 | -7.2 | -6.8 | -7.8 | -7.7 | -7.5 | -7.4 |
| CID139030492 | -8.4 | -8.2 | -7.6 | -8.2 | -9.4 | -8.36 |
| CID139030493 | -8.2 | -8.9 | -7.8 | -8 | -6.9 | -7.96 |
| CID145946018 | -8.2 | -7.4 | -7.7 | -8.1 | -7.3 | -7.74 |
| CID139030495 | -9.3 | -8.5 | -10.2 | -8.7 | -7.4 | -8.82 |
| CID139030496 | -9.3 | -8.4 | -9.9 | -8.5 | -7.3 | -8.68 |
| CID139030497 | -8.6 | -8.5 | -10.3 | -8.7 | -7.5 | -8.72 |
| CID145997887 | -9.4 | -8.4 | -9.4 | -8.7 | -10.6 | -9.3 |
| CID139030498 | -8 | -8.7 | -8.4 | -8.2 | -8.2 | -8.3 |
| CID139030500 | -9.4 | -8.5 | -8 | -8.8 | -7.7 | -8.48 |
| CID139030501 | -6.2 | -7.1 | -7.8 | -6.6 | -8.9 | -7.32 |
| CID68910199 | -6.3 | -6.7 | -7.5 | -6.4 | -8.7 | -7.12 |
| CID139030502 | -6.9 | -7 | -6.6 | -7 | -8.7 | -7.24 |
| CID71677803 | -7.7 | -7.8 | -8.7 | -8.1 | -6.9 | -7.84 |
| CID71677804 | -7.8 | -6.9 | -6.6 | -7.6 | -6.8 | -7.14 |
| CID215639 | -6.5 | -7.5 | -6.5 | -7.1 | -9 | -7.32 |
| CID56703871 | -7.2 | -8.3 | -8.5 | -7.3 | -11.2 | -8.5 |
| CID56900216 | -6.6 | -7.2 | -6.8 | -6.7 | -10.1 | -7.48 |
| CID23306 | -7.5 | -7 | -7.4 | -7.8 | -7.7 | -7.48 |
| CID4501 | -7.9 | -8.2 | -7.9 | -8 | -8.7 | -8.14 |
| CID3025944 | -6.6 | -7.2 | -8 | -7.5 | -9.2 | -7.7 |
| CID65482 | -7.2 | -8.3 | -7.5 | -7.6 | -9.3 | -7.98 |

| | Binding energy (Kcal/mol) | | | | | |
|---|---|---|---|---|---|---|
| **Pubchem CID No.** | **METTL1** | **METTL3-14** | **METTL6** | **METTL16** | **METTL18** | **Average** |
| CID5567 | -7.3 | -8 | -9.7 | -8.9 | -11.8 | -9.14 |
| CID5494506 | -9.2 | -10.7 | -9.6 | -9.1 | -7.3 | -9.18 |
| CID702558 | -7.5 | -9.2 | -8.6 | -8 | -8.2 | -8.3 |
| CID9444 | -6.5 | -6.9 | -6.4 | -6.5 | -8.2 | -6.9 |
| CID451668 | -6.3 | -6.7 | -6.4 | -6.2 | -7.8 | -6.68 |
| CID100016 | -5.9 | -6.6 | -6.5 | -6.6 | -8 | -6.72 |
| CID4914 | -5.2 | -6.6 | -6.2 | -6 | -5.4 | -5.88 |
| CID65064 | -9.2 | -8 | -9.2 | -9.1 | -7.8 | -8.66 |
| CID6400741 | -6.7 | -7 | -6.3 | -6.9 | -7.4 | -6.86 |
| CID344265 | -8.6 | -9.2 | -8.6 | -8.1 | -11.5 | -9.2 |
| CID327606 | -7.7 | -8.5 | -8.3 | -7.7 | -10.9 | -8.62 |
| CID5270713 | -5.9 | -6.7 | -6 | -6.5 | -7.4 | -6.5 |
| CID132233544 | -7.2 | -7.3 | -7.2 | -6.6 | -9.1 | -7.48 |
| CID3637 | -6.1 | -7.2 | -6.9 | -7.2 | -9.4 | -7.36 |
| CID25150857 | -9.1 | -7.8 | -9.1 | -9 | -7.5 | -8.5 |
| CID442757 | -7.1 | -8 | -7.7 | -7.8 | -7.9 | -7.7 |
| CID47751 | -5.9 | -6.8 | -6 | -6.6 | -7.4 | -6.54 |
| CID135564655 | | | | | | 0 |
| CID10037425 | -5.7 | -6.1 | -6 | -6.5 | -7.5 | -6.36 |
| CID515328 | -6.3 | -6.8 | -6.3 | -6.3 | -8.4 | -6.82 |
| CID24858111 | -9.3 | -8.8 | -12.1 | -9.4 | -8.5 | -9.62 |
| CID76318201 | -10.6 | -10.1 | -12.6 | -10.6 | -8.7 | -10.52 |
| CID156587340 | -8.2 | -8.8 | -9.9 | -9.1 | -7.3 | -8.66 |
| CID11271909 | -7.8 | -7.1 | -8.8 | -8.4 | -7.3 | -7.88 |
| CID135449332 | -10.7 | -10 | -10.7 | -9.3 | -8.1 | -9.76 |
| CID5280343 | -8.2 | -8.6 | -10.6 | -8.5 | -10.6 | -9.3 |
| CID155167581 | -9.4 | -10 | -10.1 | -9 | -10.3 | -9.76 |
| CID154815692 | -7.9 | -8.4 | -8.9 | -9.5 | -8.1 | -8.56 |
| CID60961 | -7.3 | -7.3 | -6.6 | -6.8 | -9.1 | -7.42 |
| CID98567 | -6.9 | -7.2 | -7 | -7.2 | -9.1 | -7.48 |
| CID97692 | -7.7 | -7.1 | -6.8 | -7.5 | -8.9 | -7.6 |
| CID110145076 | -8 | -7.9 | -8.2 | -7.7 | -8.9 | -8.14 |
| CID110154059 | -7.9 | -9.2 | -8.2 | -8.3 | -9.7 | -8.66 |
| CID145994381 | -7.4 | -7.2 | -7.1 | -6.8 | -8.9 | -7.48 |
| CID440041 | -5.8 | -6 | -6.2 | -5.9 | -7.4 | -6.26 |
| CID156587337 | -8.3 | -8.4 | -8 | -9.9 | -8.1 | -8.54 |
| S-Adenosyl Methionine | -7.5 | -7.7 | -7.7 | -7.3 | -10.5 | -8.14 |

Here, METTL1: Methyltransferase-like 1, METTL3: Methyltransferase-like 3, METTL6: Methyltransferase-like 6, METTL16: Methyltransferase-like 16, METTL1: Methyltransferase-like 18.

**S.table 4. DFT analysis of top 12 compounds from molecular docking analysis**

| Pubchem CID | HOMO (eV) | LUMO (eV) | Band gap eV | Hardness, Ƞ | Softness | Chemical Potential, µ |
|---|---|---|---|---|---|---|
| CID135403798 | -5.148788471 | -1.974636966 | 3.17 | 1.59 | 0.63 | -3.56 |
| CID139030487 | -4.9072 | -1.9906 | 2.92 | 1.46 | 0.69 | -3.45 |
| CID139030488 | -4.952293507 | -0.836974393 | 4.12 | 2.06 | 0.49 | -2.89 |
| CID145997887 | -5.61971242 | -1.050014009 | 4.57 | 2.28 | 0.44 | -3.33 |
| CID5567 | -6.222177244 | -2.032978636 | 4.19 | 2.09 | 0.48 | -4.13 |
| CID5494506 | -5.849541594 | -2.242426321 | 3.61 | 1.80 | 0.55 | -4.05 |
| CID344265 | -5.540581088 | -2.38866146 | 3.15 | 1.58 | 0.63 | -3.96 |
| CID24858111 | -5.332847733 | -1.97164369 | 3.36 | 1.68 | 0.60 | -3.65 |
| CID76318201 | -5.58109916 | -2.064652938 | 3.52 | 1.76 | 0.57 | -3.82 |
| CID135449332 | -5.351433256 | -1.89052591 | 3.46 | 1.73 | 0.58 | -3.62 |
| CID5280343 | -5.555955642 | -1.715392052 | 3.84 | 1.92 | 0.52 | -3.64 |
| CID155167581 | -5.4003 | -2.345 | 3.06 | 1.53 | 0.65 | -3.87 |

### S.table 5. Molecular docking results of selected 483 ZINC compounds

| | Binding energy (Kcal/mol) | | | | | | Binding energy (Kcal/mol) | | | | |
|---|---|---|---|---|---|---|---|---|---|---|---|
| ZINC ID | METTL 1 | METTL 3 | METTL 6 | METTL 16 | METTL 18 | ZINC ID | METTL 1 | METTL 3 | METTL 6 | METTL 16 | METTL 18 |
| ZINC24842130 | -8.1 | -10.7 | -8.5 | -8.8 | -11.7 | ZINC66351941 | -8.8 | -8.7 | -8.6 | -8.8 | -8 |
| ZINC03326049 | -7.7 | -8.7 | -8.2 | -8.5 | -12.4 | ZINC91057193 | -8.7 | -7.5 | -9.4 | -8.8 | -7.7 |
| ZINC36686306 | -8.6 | -8.4 | -8.5 | -8.4 | -9.2 | ZINC08093792 | -8.4 | -7.6 | -9.5 | -8.6 | -7.7 |
| ZINC03649614 | -8.4 | -8.3 | -7.2 | -8 | -10.1 | ZINC12541378 | -9 | -8 | -8.7 | -8.9 | -7.8 |
| ZINC48374334 | -8.5 | -9 | -9.1 | -8.7 | -10.3 | ZINC77099401 | -9.8 | -10.4 | -8.2 | -9.8 | -7.9 |
| ZINC23873073 | -9.3 | -8.4 | -8 | -8.4 | -11.7 | ZINC09378712 | -8.6 | -8.9 | -8.7 | -8.8 | -7.9 |
| ZINC11147774 | -8.1 | -8.2 | -10.2 | -7.9 | -11.5 | ZINC71279190 | -8.7 | -8.6 | -9.2 | -8.9 | -7.4 |
| ZINC74161832 | -8.2 | -7.5 | -9.9 | -7.8 | -10.2 | ZINC12785966 | -8.8 | -8.5 | -8.8 | -8.8 | -7.8 |
| ZINC24144770 | -8.5 | -7.8 | -10.4 | -8.2 | -11.6 | ZINC72326252 | -8.7 | -7.5 | -8.2 | -8.3 | -12.2 |
| ZINC07736825 | -8 | -7.6 | -8.7 | -7.7 | -7.5 | ZINC71285359 | -9.2 | -8.7 | -8.8 | -8.6 | -7.5 |
| ZINC21726880 | -8.6 | -9 | -9.1 | -8.6 | -7.4 | ZINC90185435 | -9 | -8.6 | -8.2 | -9.9 | -7.7 |
| ZINC15771247 | -7.5 | -7.3 | -7 | -7.4 | -9.5 | ZINC72402002 | -8.2 | -8.4 | -9.4 | -8 | -7.4 |
| ZINC12377057 | -8.4 | -8.6 | -7 | -7.6 | -6.8 | ZINC08278833 | -8.7 | -9 | -7.9 | -8.6 | -7.1 |
| ZINC81041711 | -8.4 | -8.6 | -9.3 | -8.5 | -7.4 | ZINC72401875 | -8.3 | -7.5 | -9 | -8.3 | -8.1 |
| ZINC18816560 | -8.3 | -8.1 | -9.9 | -8.4 | -10.5 | ZINC72401668 | -8.3 | -7.2 | -7.9 | -8.4 | -7.7 |
| ZINC08641984 | -7.9 | -8.2 | -7.1 | -8.6 | -10.4 | ZINC72402262 | -7.7 | -8.1 | -9.3 | -8.3 | -10.5 |
| ZINC24383880 | -7.9 | -7.8 | -9.9 | -8 | -7.4 | ZINC67216255 | -8.4 | -7.6 | -8.4 | -8.3 | -10.4 |
| ZINC19854041 | -8.6 | -8.6 | -9.5 | -8.9 | -11.4 | ZINC67216776 | -8.6 | -9.7 | -9.9 | -8.9 | -12 |
| ZINC03635872 | -8.9 | -8.9 | -8.8 | -8.9 | -11.3 | ZINC67216544 | -8.6 | -8.9 | -10.4 | -9 | -12.4 |
| ZINC09858672 | -8.5 | -8 | -8 | -8 | -11.3 | ZINC91443590 | -9.5 | -8.6 | -8.3 | -9.3 | -7.5 |
| ZINC16321499 | -8 | -9.9 | -9.1 | -8.3 | -7.2 | ZINC67217744 | -9.3 | -8.2 | -8.5 | -8.4 | -11.9 |
| ZINC09845394 | -8.9 | -10.2 | -10.5 | -10.2 | -11.1 | ZINC12985894 | -8.6 | -8.8 | -9.9 | -9.1 | -11.8 |
| ZINC09845625 | -8.1 | -7.4 | -8 | -7.7 | -10.3 | ZINC67216666 | -9.2 | -8.5 | -9.3 | -9.2 | -7.8 |
| ZINC21284832 | -7.3 | -6.7 | -8.5 | -7.3 | -7 | ZINC77505524 | -8.5 | -9.2 | -9.2 | -8.2 | -7.8 |
| ZINC10367096 | -7.2 | -7.2 | -6.3 | -7.2 | -7 | ZINC40039977 | -9.4 | -10.6 | -11 | -9.7 | -11.7 |
| ZINC03542773 | -8.6 | -8.3 | -9.7 | -7.9 | -11.1 | ZINC33246308 | -9.2 | -9 | -11 | -10.3 | -11.8 |
| ZINC23845829 | -8.7 | -8.8 | -9.4 | -9 | -11.2 | ZINC49609445 | -8.8 | -9.5 | -8.3 | -9 | -8.1 |
| ZINC13688653 | -8.3 | -7.8 | -9.1 | -7.9 | -10.5 | ZINC61234599 | -8.9 | -7.7 | -8.9 | -8 | -10.4 |
| ZINC07214993 | -8.4 | -7.8 | -8.7 | -7.6 | -9.2 | ZINC49625242 | -8.8 | -8.2 | -8.8 | -8.4 | -12.2 |
| ZINC09955798 | -8.7 | -7.5 | -7.4 | -8.1 | -10.4 | ZINC33594885 | -7.3 | -7.9 | -7.6 | -7.9 | -7.4 |
| ZINC89914794 | -8.3 | -9.5 | -7.3 | -7.4 | -11.1 | ZINC54466629 | -8.6 | -7.7 | -8.1 | -8.1 | -7.8 |
| ZINC12390177 | -8.3 | -8.6 | -9.1 | -8.8 | -7.5 | ZINC28690681 | -8.6 | -7.9 | -7.8 | -7.7 | -10.6 |
| ZINC12115283 | -8.5 | -8.3 | -7.9 | -8.3 | -10.2 | ZINC28714964 | -8.7 | -7.5 | -7.9 | -7.5 | -8.2 |
| ZINC85740282 | -8 | -8.4 | -7.9 | -8.7 | -8.2 | ZINC28715060 | -9.7 | -8.9 | -9.4 | -8.8 | -8.1 |
| ZINC94125143 | -8.2 | -7.7 | -7.8 | -8.3 | -10.9 | ZINC72410304 | -9.6 | -8.5 | -9.4 | -8.7 | -8.1 |
| ZINC32926152 | -8.2 | -8 | -8.8 | -7.9 | -10.1 | ZINC12279561 | -8.8 | -9.1 | -8.5 | -8.9 | -7.3 |
| ZINC72336185 | -8.4 | -9.5 | -7.3 | -7.8 | -7.2 | ZINC28714806 | -9.6 | -9.1 | -9.1 | -8.7 | -7.9 |
| ZINC94127054 | -8.1 | -8 | -8.5 | -8.4 | -7.1 | ZINC08642836 | -9.6 | -8.8 | -8.6 | -8.6 | -8.1 |
| ZINC72336186 | -8.2 | -8.4 | -8.8 | -8.5 | -9.5 | ZINC14979676 | -8.3 | -8.2 | -7 | -8.9 | -10.4 |
| ZINC94013431 | -7.8 | -8 | -8.5 | -7.7 | -9.7 | ZINC12765331 | -9.4 | -8.5 | -11.4 | -9.1 | -7.9 |
| ZINC32670106 | -8.1 | -8 | -7.4 | -8.4 | -10 | ZINC03333248 | -8.9 | -8.5 | -8.9 | -8.5 | -11.8 |

| ZINC ID | Binding energy (kcal/mol) | | | | | ZINC ID | Binding energy (kcal/mol) | | | | |
|---|---|---|---|---|---|---|---|---|---|---|---|
| | METTL 1 | METTL 3 | METTL 6 | METTL 16 | METTL 18 | | METTL 1 | METTL 3 | METTL 6 | METTL 16 | METTL 18 |
| ZINC72280514 | -8.6 | -10.1 | -8.4 | -8.4 | -8 | ZINC17087123 | -8.6 | -9 | -9.1 | -9.3 | -12.1 |
| ZINC23805266 | -8 | -8.4 | -9.4 | -8.5 | -11.8 | ZINC23632079 | -9.2 | -10.7 | -9.8 | -9.2 | -11.9 |
| ZINC13963680 | -8.2 | -9.2 | -7.4 | -8.8 | -11.2 | ZINC71284713 | -8.7 | -11.8 | -9.9 | -8.9 | -7.7 |
| ZINC23883179 | -8.2 | -8.6 | -7.9 | -8.2 | -7.4 | ZINC92860505 | -8.9 | -7.7 | -8.2 | -8.2 | -7.5 |
| ZINC23989898 | -8.2 | -7.8 | -7 | -8.4 | -10.4 | ZINC36580060 | -9 | -10.5 | -9.2 | -8.8 | -11.7 |
| ZINC23883274 | -8.2 | -8.5 | -7.1 | -8.5 | -7.4 | ZINC02257180 | -9.4 | -8.8 | -10 | -8.5 | -11.4 |
| ZINC11248784 | -7.4 | -6.7 | -9 | -7 | -10 | ZINC02255478 | -9.1 | -7.6 | -8 | -8.2 | -11.4 |
| ZINC11663767 | -7.4 | -7.4 | -6.5 | -7 | -6.8 | ZINC02256741 | -8.8 | -7.9 | -8.3 | -9 | -12.8 |
| ZINC14244578 | -8.1 | -9 | -7.2 | -7.9 | -7 | ZINC02247247 | -8.7 | -9.8 | -7.8 | -8.1 | -11.2 |
| ZINC25184885 | -8 | -8.2 | -8.3 | -8.2 | -10.3 | ZINC02253645 | -7.8 | -8.1 | -7 | -7.8 | -10.1 |
| ZINC10587047 | -8.8 | -7.9 | -9 | -8.5 | -11.6 | ZINC70666323 | -8.3 | -9.8 | -8.4 | -7.4 | -11.1 |
| ZINC03671877 | -8.5 | -10.2 | -10.3 | -8.6 | -7.7 | ZINC61721532 | -8.8 | -10.4 | -8 | -8.6 | -7.7 |
| ZINC74162504 | -7.9 | -9.8 | -7.3 | -7.7 | -8.1 | ZINC70666503 | -9.4 | -10.4 | -11.1 | -9.1 | -13.1 |
| ZINC12261832 | -8.6 | -9.9 | -8.6 | -8.3 | -12.1 | ZINC72401166 | -8.7 | -8.3 | -8.7 | -8.3 | -8 |
| ZINC28628873 | -7.7 | -7.5 | -7 | -7.4 | -7.1 | ZINC14180608 | -8.1 | -8.3 | -9 | -8.5 | -7.2 |
| ZINC36800408 | -8.4 | -8.6 | -8.3 | -8.2 | -7.8 | ZINC40324131 | -8.2 | -9.1 | -8.4 | -8.2 | -7.4 |
| ZINC14370707 | -7.6 | -6.9 | -8.3 | -7.7 | -10.2 | ZINC72401056 | -9.7 | -9.6 | -9.6 | -8.8 | -8.6 |
| ZINC20866616 | -7.9 | -7.3 | -7.6 | -8.2 | -9.8 | ZINC11951787 | -8 | -9.3 | -8.4 | -8 | -7.1 |
| ZINC16431285 | -8.1 | -8.4 | -9.4 | -8.7 | -10.4 | ZINC69260921 | -9.1 | -7.1 | -7.8 | -8.7 | -12.4 |
| ZINC20866661 | -8.1 | -7.6 | -8.3 | -8.7 | -10.4 | ZINC40324129 | -8 | -7.4 | -9 | -8.4 | -7.4 |
| ZINC94130164 | -8.4 | -10.3 | -9.6 | -8.4 | -7.8 | ZINC40324133 | -9.4 | -8.6 | -8.2 | -8.9 | -12.2 |
| ZINC94018200 | -9 | -10 | -8.7 | -8.9 | -11.7 | ZINC36621797 | -8.9 | -9.4 | -8.9 | -8.4 | -11.7 |
| ZINC90559514 | -8.4 | -9.4 | -7.8 | -8.1 | -7.9 | ZINC72302744 | -8.1 | -10.8 | -7.6 | -9 | -12.4 |
| ZINC05396560 | -8.6 | -8.8 | -10.3 | -9.2 | -8.1 | ZINC40324139 | -9.4 | -8.1 | -9.9 | -9.2 | -8.1 |
| ZINC09507128 | -8.6 | -8.1 | -8.7 | -9 | -11.6 | ZINC95360112 | -8 | -8.1 | -7.3 | -7.8 | -7.3 |
| ZINC07313162 | -8.3 | -7 | -7.7 | -8.4 | -7.4 | ZINC09243441 | -7.4 | -8.9 | -7.1 | -9.1 | -6.5 |
| ZINC03405008 | -7.9 | -8.6 | -8.1 | -8.1 | -7.3 | ZINC89494973 | -8.9 | -8.4 | -7.8 | -9.3 | -7.8 |
| ZINC14910357 | -8.2 | -7.8 | -9.5 | -8.3 | -7.5 | ZINC09124297 | -9.4 | -8.9 | -11.1 | -9.2 | -11.9 |
| ZINC93906186 | -8.6 | -8.3 | -9.1 | -8.6 | -10.9 | ZINC33256561 | -8.7 | -9.2 | -9.8 | -7.9 | -7.8 |
| ZINC31505912 | -8.8 | -8.2 | -8.3 | -8.9 | -11.5 | ZINC20415103 | -9.2 | -8.7 | -8.4 | -8.4 | -11.9 |
| ZINC12340933 | -9.1 | -8.3 | -7.8 | -8.7 | -6.9 | ZINC72403395 | -8.9 | -7.4 | -7.9 | -8.4 | -10.8 |
| ZINC90642692 | -9.4 | -10.2 | -9.5 | -10.1 | -7.9 | ZINC81795442 | -9.2 | -8.1 | -9.6 | -9.2 | -8.3 |
| ZINC07313163 | -8.6 | -9.8 | -7.5 | -7.9 | -12.4 | ZINC40762990 | -8.9 | -9.7 | -9.8 | -9.2 | -7.9 |
| ZINC10320175 | -9.5 | -8.3 | -8.6 | -9.9 | -8 | ZINC05938302 | -8.3 | -7.1 | -8.8 | -8.2 | -9.2 |
| ZINC23873131 | -7.5 | -7.2 | -6.2 | -6.6 | -10.3 | ZINC70931093 | -9.1 | -8.9 | -9.1 | -8.9 | -8.3 |
| ZINC85875346 | -7.3 | -6.8 | -6.6 | -7.1 | -10 | ZINC79153644 | -9.3 | -8.1 | -8.5 | -9 | -8 |
| ZINC94099789 | -8.6 | -10.1 | -8.4 | -8.2 | -12.5 | ZINC91033029 | -8.1 | -8.1 | -8.9 | -8.2 | -8.1 |
| ZINC93909723 | -8.3 | -8.1 | -8.9 | -9.3 | -12.2 | ZINC10690826 | -8.1 | -7.3 | -8.8 | -8 | -9.1 |
| ZINC12530383 | -8.7 | -8.9 | -8.7 | -8.4 | -7.8 | ZINC12575148 | -8.8 | -7.6 | -7.8 | -8.1 | -8.5 |
| ZINC72024370 | -8.2 | -8 | -8.8 | -9.7 | -7.9 | ZINC25139533 | -9 | -8.7 | -9.2 | -9.4 | -10.7 |
| ZINC08093793 | -8.3 | -9.1 | -8.6 | -10 | -7.8 | ZINC13000658 | -9.5 | -9.8 | -10 | -9 | -11.8 |
| ZINC03427622 | -8.4 | -8.2 | -7.4 | -8.6 | -7.2 | ZINC36801391 | -9.4 | -9.9 | -10.6 | -9.6 | -8.3 |

| ZINC ID | Binding energy (kcal/mol) | | | | | ZINC ID | Binding energy (kcal/mol) | | | | |
|---|---|---|---|---|---|---|---|---|---|---|---|
| | METTL 1 | METTL 3 | METTL 6 | METTL 16 | METTL 18 | | METTL 1 | METTL 3 | METTL 6 | METTL 16 | METTL 18 |
| ZINC74273421 | -8.4 | -8.7 | -9.1 | -9.3 | -12.7 | ZINC06984862 | -8.5 | -8.9 | -9.7 | -9.8 | -8 |
| ZINC90644988 | -8.3 | -9.4 | -9.6 | -8.8 | -8 | ZINC70931344 | -8.8 | -8.8 | -8.1 | -9 | -8 |
| ZINC07987235 | -7.6 | -10.3 | -8.8 | -9.4 | -12 | ZINC40049280 | -8.9 | -8.1 | -9.9 | -7.9 | -7.8 |
| ZINC04493720 | -8.5 | -8.9 | -9.7 | -8.5 | -12 | ZINC78920057 | -8.5 | -9 | -7.9 | -9.2 | -7.3 |
| ZINC12793212 | -7.7 | -8.7 | -7.9 | -9.1 | -12.5 | ZINC57868915 | -8.3 | -8.1 | -8.1 | -8.5 | -7.9 |
| ZINC59268385 | -8.5 | -8.1 | -9.3 | -8.3 | -12.1 | ZINC03406473 | -8.8 | -7.7 | -8.4 | -8.1 | -7.9 |
| ZINC90643103 | -8.9 | -9.9 | -9.4 | -8.4 | -10.3 | ZINC67794556 | -9.2 | -8.4 | -8.2 | -8.8 | -11.9 |
| ZINC67217484 | -9.1 | -10 | -9 | -8.7 | -7.4 | ZINC81649707 | -9.8 | -10.2 | -8.2 | -9.5 | -8.3 |
| ZINC33941516 | -8.2 | -8.7 | -8.5 | -9 | -7.7 | ZINC72242996 | -9.2 | -8.7 | -10.1 | -9.6 | -8.7 |
| ZINC06198009 | -8.5 | -8.7 | -9.7 | -8.5 | -11.5 | ZINC08429769 | -10.2 | -10.3 | -10.2 | -9.9 | -8.4 |
| ZINC08039700 | -8.6 | -8.6 | -9 | -8.8 | -7.4 | ZINC12276217 | -9.6 | -9.4 | -9.7 | -9.4 | -7.9 |
| ZINC06536284 | -8.8 | -9.5 | -7.6 | -8.8 | -10.9 | ZINC12360516 | -7.8 | -9.8 | -7.3 | -9.2 | -12.1 |
| ZINC02788826 | -8.1 | -8 | -7.5 | -8.9 | -10.8 | ZINC19563153 | -7.8 | -7.9 | -7.6 | -9.1 | -11.8 |
| ZINC91987006 | -8 | -8.7 | -9.2 | -8.6 | -7.2 | ZINC18948564 | -8.8 | -8.3 | -8.6 | -9.7 | -7.8 |
| ZINC21726936 | -8.9 | -7.9 | -7.9 | -8.4 | -9.2 | ZINC40050977 | -9.1 | -8.6 | -9.6 | -8.9 | -11.7 |
| ZINC44120706 | -8.5 | -8.6 | -8.5 | -8.3 | -10.4 | ZINC91074718 | -8.9 | -8.7 | -8.6 | -9.4 | -7.4 |
| ZINC33328929 | -8.8 | -8.4 | -8.4 | -8.2 | -10.6 | ZINC12506923 | -7.7 | -8.3 | -6.6 | -7.9 | -9.1 |
| ZINC15868040 | -8 | -8.2 | -8.9 | -8.2 | -10.5 | ZINC91073191 | -8.8 | -7.4 | -7.9 | -8.7 | -8 |
| ZINC79691506 | -8.7 | -8.5 | -8.7 | -8.8 | -11 | ZINC66473853 | -8.6 | -7.4 | -7.8 | -8.4 | -8.2 |
| ZINC58435734 | -8.1 | -8.7 | -8.4 | -8.2 | -7.7 | ZINC54466355 | -9.2 | -8.2 | -9.4 | -8.9 | -7.9 |
| ZINC94068856 | -8.3 | -8.6 | -7.9 | -7.9 | -9.7 | ZINC05332359 | -9.4 | -8.5 | -10.2 | -10 | -8.3 |
| ZINC94157554 | -8.2 | -8.7 | -7.9 | -9.6 | -7.8 | ZINC91074745 | -8.2 | -7 | -8.7 | -7.7 | -7.8 |
| ZINC19117040 | -7.5 | -9.5 | -9.6 | -8.5 | -12.3 | ZINC10546698 | -9.3 | -10.9 | -10.4 | -10.6 | -7.7 |
| ZINC71885476 | -8 | -10 | -7.7 | -8.2 | -7.1 | ZINC91065910 | -6.9 | -8.6 | -6.5 | -9.3 | -8 |
| ZINC58177766 | -9 | -8 | -8.2 | -8.3 | -11.3 | ZINC62588334 | -8.2 | -8.8 | -9.6 | -8.8 | -7.4 |
| ZINC90644447 | -8.6 | -8.2 | -8.3 | -9 | -11.1 | ZINC33252787 | -9.2 | -8.3 | -8.8 | -10.1 | -13.1 |
| ZINC95358659 | -8.5 | -7.9 | -7.4 | -8.2 | -12.6 | ZINC67935470 | -8.4 | -8.7 | -8.1 | -8.1 | -12.5 |
| ZINC02782188 | -7.5 | -8.3 | -7.7 | -8.4 | -7.3 | ZINC31782528 | -8.9 | -8.6 | -9.8 | -9.2 | -8 |
| ZINC04121341 | -8.8 | -7.8 | -7.8 | -7.8 | -7.3 | ZINC49414161 | -9.4 | -8.8 | -8.2 | -9.2 | -8.6 |
| ZINC38600495 | -7.5 | -7.1 | -9.2 | -8.6 | -11.9 | ZINC06198307 | -8.8 | -8 | -7.7 | -8.4 | -11.4 |
| ZINC02795221 | -8.3 | -8.6 | -10.5 | -8.5 | -11.4 | ZINC06829939 | -8.5 | -8.8 | -8.5 | -9.2 | -8 |
| ZINC08278840 | -8.1 | -8 | -10 | -8.1 | -7.4 | ZINC49600780 | -9.8 | -8.6 | -9.9 | -9.1 | -8.1 |
| ZINC01009161 | -9 | -9.9 | -8.6 | -8.5 | -12.2 | ZINC93913479 | -8.5 | -8.7 | -6.5 | -8.7 | -12 |
| ZINC89204531 | -8.5 | -8.8 | -9.5 | -8.9 | -7.8 | ZINC19720350 | -9.1 | -10.1 | -10.3 | -9.4 | -10.2 |
| ZINC02778927 | -8.6 | -8.8 | -7.5 | -9.2 | -11.8 | ZINC91452445 | -8.5 | -8.2 | -7.3 | -8.3 | -11 |
| ZINC40071682 | -8.3 | -8.2 | -9 | -8 | -7.2 | ZINC09379525 | -8.3 | -7.5 | -7.3 | -8.4 | -7.7 |
| ZINC17011809 | -8.8 | -8.5 | -10.7 | -9.2 | -12.6 | ZINC71285389 | -9 | -8.3 | -10.5 | -9.8 | -8.5 |
| ZINC25249143 | -8.3 | -10.8 | -10.7 | -9.1 | -13 | ZINC39372372 | -8.8 | -9.7 | -9.2 | -8.7 | -9.9 |
| ZINC02887051 | -8.7 | -8.1 | -8.6 | -9 | -8.1 | ZINC18203558 | -8.9 | -9.7 | -9.5 | -9.2 | -8 |
| ZINC60387170 | -8.9 | -9.9 | -8.5 | -9.9 | -11.7 | ZINC02458551 | -8.5 | -8.7 | -8.8 | -9.4 | -12.9 |
| ZINC20442293 | -8.9 | -9.1 | -8.2 | -9.5 | -11.9 | ZINC06383185 | -7.9 | -9.2 | -8.3 | -9.9 | -13.7 |
| ZINC40661673 | -8.4 | -8.6 | -9.6 | -8.2 | -11.1 | ZINC40762889 | -8.3 | -7.7 | -7.9 | -8 | -7.9 |

| ZINC ID | Binding energy (kcal/mol) | | | | | ZINC ID | Binding energy (kcal/mol) | | | | |
|---|---|---|---|---|---|---|---|---|---|---|---|
| | METTL 1 | METTL 3 | METTL 6 | METTL 16 | METTL 18 | | METTL 1 | METTL 3 | METTL 6 | METTL 16 | METTL 18 |
| ZINC90643983 | -8.3 | -9 | -9.1 | -8.6 | -7 | ZINC30972905 | -9.2 | -9.3 | -8.4 | -8.8 | -9.2 |
| ZINC89869338 | -7.2 | -7.4 | -6.7 | -6.9 | -7.3 | ZINC39916723 | -8.4 | -8.5 | -8.4 | -8.5 | -7.8 |
| ZINC90530846 | -9 | -8.4 | -8 | -9.4 | -11.8 | ZINC22201396 | -7.6 | -6.7 | -7.2 | -7.2 | -7.4 |
| ZINC17172072 | -7.8 | -7.5 | -8.3 | -8.4 | -10.1 | ZINC40759363 | -8.6 | -8.2 | -7.6 | -8.8 | -11.5 |
| ZINC55396116 | -8 | -10.1 | -9.4 | -8 | -11.7 | ZINC64372616 | -8.6 | -8.2 | -9.4 | -7.9 | -7.9 |
| ZINC79692364 | -8.7 | -7.8 | -6.5 | -7.3 | -10.1 | ZINC12445585 | -9.2 | -8.6 | -10.1 | -9.3 | -8.1 |
| ZINC15601815 | -7.8 | -7.9 | -7.4 | -8.3 | -12.1 | ZINC78778593 | -8.6 | -9 | -9.6 | -10 | -8.1 |
| ZINC90644974 | -9.1 | -9 | -7.9 | -8 | -12.2 | ZINC39916724 | -8.8 | -8.1 | -9.4 | -9 | -7.8 |
| ZINC90643097 | -7.8 | -8.8 | -7.8 | -9 | -12.1 | ZINC40760137 | -10.1 | -9.2 | -10.7 | -9.2 | -8.2 |
| ZINC90644976 | -8.6 | -8.6 | -8.7 | -8.1 | -10.4 | ZINC40759749 | -8.7 | -8 | -10.2 | -8.1 | -7.7 |
| ZINC90643099 | -8.9 | -8.5 | -10.1 | -8.4 | -8.1 | ZINC40758977 | -8.9 | -8.2 | -9.4 | -9 | -8.1 |
| ZINC22473005 | -8.6 | -8.4 | -8.8 | -7.9 | -7.1 | ZINC04371075 | -10 | -9.8 | -8.9 | -9.9 | -8.6 |
| ZINC27795763 | -8.6 | -9.8 | -11 | -8.9 | -12.7 | ZINC40758207 | -8.6 | -8 | -7.4 | -8.6 | -10.9 |
| ZINC27793958 | -8 | -9.8 | -7.6 | -7.1 | -11.4 | ZINC77437423 | -8.3 | -6.9 | -7.5 | -8.4 | -10.8 |
| ZINC15567207 | -8.4 | -9 | -7 | -7.4 | -8.2 | ZINC31849451 | -9.1 | -7.8 | -7.5 | -7.6 | -8.3 |
| ZINC90533356 | -8.7 | -8.8 | -7.8 | -9.1 | -10.8 | ZINC24527699 | -8.7 | -8.3 | -8.6 | -9.1 | -8.8 |
| ZINC89294352 | -8.7 | -8.4 | -10 | -9.2 | -7.5 | ZINC40145196 | -9.2 | -8.6 | -8.2 | -10.2 | -8.1 |
| ZINC01499343 | -8.8 | -9.3 | -9.3 | -8.4 | -11.5 | ZINC81649853 | -8.6 | -9.3 | -8.5 | -9.5 | -8.5 |
| ZINC67794559 | -8.6 | -7.7 | -7.4 | -9 | -10.6 | ZINC32090248 | -8.1 | -8.1 | -9.8 | -9.1 | -7.8 |
| ZINC02209747 | -8.5 | -8.8 | -9 | -9 | -8.1 | ZINC23856091 | -8.4 | -9.5 | -8 | -8.7 | -12.3 |
| ZINC44120689 | -9.2 | -8.4 | -8.9 | -9 | -7.8 | ZINC72324946 | -9.2 | -8.8 | -10.8 | -9.2 | -12.3 |
| ZINC36709136 | -9.7 | -8.2 | -8.4 | -9.4 | -8.3 | ZINC95350884 | -9.5 | -9 | -9 | -9.9 | -7.3 |
| ZINC25227568 | -8.7 | -8.7 | -8.3 | -8.5 | -12 | ZINC72324978 | -8.9 | -8.5 | -8.8 | -9.6 | -8.7 |
| ZINC12149176 | -8.5 | -9 | -9 | -8.9 | -11.6 | ZINC90185427 | -9.2 | -8.7 | -9.4 | -9 | -8 |
| ZINC05241819 | -9.4 | -9.2 | -8.2 | -8.3 | -12.2 | ZINC02350676 | -8.7 | -8.2 | -7.4 | -8.9 | -7.6 |
| ZINC68785853 | -8.5 | -8.7 | -8 | -8.1 | -7.5 | ZINC92963722 | -8.8 | -8.8 | -9 | -8.4 | -6.8 |
| ZINC16755273 | -8.1 | -7.5 | -8.8 | -7.8 | -11.2 | ZINC36636549 | -9 | -8.2 | -7.9 | -9.1 | -11.6 |
| ZINC03337042 | -8 | -9.6 | -7.4 | -8.1 | -11.3 | ZINC45673942 | -8.9 | -7.9 | -7.2 | -8.6 | -7.3 |
| ZINC13958237 | -8.1 | -7.6 | -7.2 | -8.9 | -7.8 | ZINC12414137 | -8.8 | -8.1 | -8.6 | -9.2 | -8.4 |
| ZINC18699413 | -9.2 | -8.5 | -7.9 | -8.2 | -11.8 | ZINC24950359 | -8.2 | -7.2 | -7.5 | -7.4 | -9.9 |
| ZINC01032649 | -9 | -9.8 | -10.4 | -9.1 | -8.1 | ZINC11612151 | -9.2 | -9.3 | -10.2 | -9.5 | -8.2 |
| ZINC13729760 | -8.5 | -9.7 | -9 | -8.5 | -11.4 | ZINC90150043 | -7.9 | -9.8 | -7.4 | -7.6 | -7.3 |
| ZINC89420646 | -8.6 | -7.9 | -9.3 | -8.6 | -10.6 | ZINC91270381 | -9.3 | -9 | -10.2 | -9.5 | -7.8 |
| ZINC02418420 | -7.8 | -7.9 | -10 | -8 | -7.2 | ZINC27912364 | -9.1 | -9.6 | -9.1 | -8.7 | -8 |
| ZINC91073189 | -8.3 | -7 | -7.5 | -7.5 | -7.5 | ZINC32900586 | -10.2 | -9.6 | -9.4 | -8.8 | -8 |
| ZINC90644586 | -8.5 | -7.3 | -9.4 | -8.5 | -7.6 | ZINC40143006 | -9.3 | -8.7 | -8.5 | -8.7 | -8.5 |
| ZINC78772377 | -8.2 | -9.4 | -7.7 | -7.8 | -7.3 | ZINC19714372 | -9.3 | -8.3 | -8.5 | -9.5 | -11.1 |
| ZINC25426432 | -8.1 | -9.1 | -9.1 | -8.8 | -7.8 | ZINC72325479 | -8.4 | -9.6 | -9.3 | -8 | -7.6 |
| ZINC91022641 | -8.1 | -8 | -8.3 | -7.3 | -11 | ZINC07670244 | -9.2 | -8.2 | -10.2 | -9.3 | -8 |
| ZINC25747894 | -8.6 | -8.4 | -10.1 | -8.5 | -10.9 | ZINC31842698 | -9.2 | -7 | -8.2 | -8 | -9.9 |
| ZINC57701057 | -8.2 | -7.8 | -7.5 | -9.3 | -11.4 | ZINC39916727 | -8.9 | -8.1 | -9.3 | -9 | -7.6 |
| ZINC12935539 | -8.6 | -8.1 | -10.2 | -8.7 | -11.1 | ZINC39916726 | -8.7 | -8.6 | -8.2 | -9.6 | -7.4 |

| ZINC ID | Binding energy (kcal/mol) METTL 1 | METTL 3 | METTL 6 | METTL 16 | METTL 18 | ZINC ID | Binding energy (kcal/mol) METTL 1 | METTL 3 | METTL 6 | METTL 16 | METTL 18 |
|---|---|---|---|---|---|---|---|---|---|---|---|
| ZINC14891501 | -9.3 | -9.1 | -10.2 | -9.6 | -12.8 | ZINC33040094 | -9.1 | -9.3 | -9.2 | -8.6 | -7.7 |
| ZINC12585333 | -9 | -8.5 | -8.8 | -8.5 | -11.5 | ZINC02085838 | -9.1 | -8.4 | -9.5 | -9.7 | -7.3 |
| ZINC69423180 | -8.9 | -8.8 | -8.4 | -8.8 | -8 | ZINC91038283 | -9.7 | -8.8 | -8.8 | -8.6 | -8.2 |
| ZINC16430932 | -8.4 | -7.9 | -8 | -8 | -8.1 | ZINC39287646 | -9.8 | -8.5 | -9.3 | -10.4 | -8.8 |
| ZINC89868646 | -8.5 | -8.5 | -8 | -8.8 | -8 | ZINC39288371 | -8.8 | -8.3 | -9.9 | -9.6 | -7.2 |
| ZINC02802432 | -8.1 | -8.7 | -8.6 | -8.3 | -7.6 | ZINC12428067 | -9.6 | -9.4 | -11.7 | -9.6 | -13.9 |
| ZINC12534768 | -8.8 | -9 | -8.3 | -8.3 | -7.6 | ZINC39289259 | -8.5 | -7.3 | -6.6 | -7.4 | -8.7 |
| ZINC39357459 | -8.5 | -8.3 | -8.6 | -8.1 | -7.6 | ZINC19714767 | -11 | -9.3 | -9.8 | -9.5 | -8.4 |
| ZINC11879768 | -9.5 | -9.3 | -8.1 | -8.3 | -7.8 | ZINC90024715 | -8.3 | -8.9 | -8.3 | -9.4 | -7.9 |
| ZINC02802726 | -8.8 | -8.9 | -7.9 | -9 | -7.9 | ZINC10320176 | -9.2 | -10.7 | -13.3 | -11.5 | -10.9 |
| ZINC90999035 | -6.9 | -6.7 | -6.8 | -7 | -7.3 | ZINC21962741 | -9.3 | -8.5 | -10 | -8.3 | -7.8 |
| ZINC11762732 | -9.1 | -9 | -9.2 | -9.2 | -10.1 | ZINC12639638 | -7.9 | -6.3 | -7.2 | -8.3 | -6.8 |
| ZINC00720896 | -8.4 | -9.1 | -7.8 | -8.4 | -13.2 | ZINC11545563 | -8.7 | -8 | -9.2 | -8.5 | -7.7 |
| ZINC03294047 | -8.2 | -9 | -6.9 | -7.9 | -7.3 | ZINC61088915 | -9.7 | -8.1 | -7.6 | -9.1 | -7.2 |
| ZINC71959778 | -7.2 | -5.9 | -6.2 | -6.4 | -6.5 | ZINC68569859 | -9.5 | -7.9 | -8.7 | -10.3 | -8.3 |
| ZINC89851292 | -7.6 | -7.1 | -6.8 | -7.1 | -10.7 | ZINC11546338 | -9.4 | -9.2 | -9.9 | -9.4 | -9.2 |
| ZINC25747900 | -9.1 | -8.6 | -8 | -8.7 | -10.7 | ZINC11546433 | -8.2 | -8 | -7.9 | -8.9 | -7.6 |
| ZINC92860595 | -8.6 | -9.5 | -8.6 | -8.7 | -12.2 | ZINC13020837 | -9.4 | -7.9 | -7.7 | -9.6 | -12.3 |
| ZINC67734495 | -9.2 | -8.1 | -10.1 | -9.6 | -12.8 | ZINC12980524 | -8.7 | -8.8 | -8.2 | -9.9 | -7.7 |
| ZINC77420728 | -7.7 | -8.1 | -7.2 | -8 | -10 | ZINC40040596 | -9.4 | -8.4 | -9.6 | -8.8 | -8.9 |
| ZINC06759083 | -8.4 | -8.2 | -7.6 | -8.6 | -7.8 | ZINC91031735 | -9.2 | -8.5 | -8.1 | -8.9 | -7.2 |
| ZINC12664591 | -8.4 | -7.6 | -8.6 | -8 | -10.5 | ZINC08279024 | -8 | -7.8 | -7.7 | -7.6 | -8.3 |
| ZINC12664621 | -9 | -8.4 | -8.7 | -8.9 | -12.4 | ZINC42038827 | -8.8 | -8.8 | -8.2 | -8.4 | -8 |
| ZINC06028992 | -9.1 | -10.3 | -7.9 | -9.5 | -12.6 | ZINC23054928 | -9.5 | -9.3 | -9.6 | -10 | -8.6 |
| ZINC03332474 | -8.7 | -8.7 | -9 | -8.6 | -10.3 | ZINC91044858 | -8.7 | -8.1 | -8.6 | -8.3 | -7.2 |
| ZINC40312663 | -9.3 | -8.1 | -7.6 | -8.7 | -7.9 | ZINC90999060 | -9.2 | -9.6 | -8.8 | -8.9 | -10 |
| ZINC90645059 | -8.6 | -8.1 | -9.7 | -8.6 | -12.4 | ZINC91057191 | -9.8 | -8.6 | -8.5 | -10.1 | -8.6 |
| ZINC90643131 | -8.2 | -8.5 | -7.9 | -8.1 | -7.9 | ZINC09376468 | -9.3 | -8.4 | -8.6 | -9.2 | -8.2 |
| ZINC19126493 | -9.1 | -8.7 | -8 | -9 | -14.1 | ZINC91738511 | -8.2 | -7.1 | -8.1 | -8.2 | -7.2 |
| ZINC72266025 | -9.1 | -10.5 | -10.9 | -10 | -7.7 | ZINC22709430 | -7.6 | -7.6 | -7.6 | -7.8 | -7.3 |
| ZINC15065068 | -7.7 | -7.8 | -8.3 | -8.8 | -11.1 | ZINC89434048 | -9.5 | -8.6 | -8 | -8 | -7.8 |
| ZINC65477032 | -8.9 | -8.5 | -7.7 | -8.2 | -11.7 | ZINC81650061 | -8.9 | -8 | -8.3 | -9.5 | -7.8 |
| ZINC12664569 | -8.5 | -10.3 | -8 | -8.4 | -12 | ZINC12529399 | -7.2 | -8 | -7.2 | -7.5 | -7.1 |
| ZINC40309820 | -8.6 | -10.4 | -8.2 | -8.5 | -12 | ZINC92914960 | -9.2 | -8.5 | -8 | -8.9 | -8.3 |
| ZINC58433601 | -8.7 | -7.9 | -8.3 | -9.2 | -11.2 | ZINC90644742 | -9.2 | -9 | -8.3 | -8.3 | -7.6 |
| ZINC20332294 | -8.7 | -7.8 | -8.1 | -8.7 | -11 | ZINC90642973 | -9.4 | -9 | -9.6 | -9.7 | -8.3 |
| ZINC11150819 | -8 | -9.1 | -7.7 | -8.3 | -11.1 | ZINC92922979 | -8.3 | -11 | -10.6 | -8.9 | -7.6 |
| ZINC90642862 | -9.2 | -10.4 | -11 | -10.1 | -7.8 | ZINC68569709 | -8.9 | -8.9 | -8.8 | -9 | -8 |
| ZINC95368362 | -8.2 | -9 | -8.1 | -8.7 | -11.7 | ZINC40068901 | -8.6 | -7.6 | -8 | -10.3 | -8.5 |
| ZINC69607343 | -9.3 | -9.6 | -9.7 | -9 | -7.8 | ZINC90644007 | -9.4 | -8.1 | -8.2 | -8.8 | -7.7 |
| ZINC77437424 | -9.2 | -8.4 | -9.3 | -8.3 | -7.7 | ZINC90642390 | -9.8 | -10 | -8.2 | -9.5 | -7.4 |
| ZINC32961253 | -8.9 | -8.2 | -8.9 | -9 | -7.8 | ZINC04121318 | -8.8 | -9.2 | -8.7 | -9.1 | -8.5 |

| ZINC ID | Binding energy (kcal/mol) METTL 1 | METTL 3 | METTL 6 | METTL 16 | METTL 18 | ZINC ID | Binding energy (kcal/mol) METTL 1 | METTL 3 | METTL 6 | METTL 16 | METTL 18 |
|---|---|---|---|---|---|---|---|---|---|---|---|
| ZINC05353138 | -7 | -9.2 | -6.7 | -8.4 | -6 | ZINC12891381 | -9 | -11.6 | -9 | -9.5 | -8.1 |
| ZINC32961116 | -8.5 | -7.9 | -7.4 | -7.8 | -11.7 | ZINC11936519 | -9.1 | -9.3 | -10.1 | -9.3 | -7.9 |
| ZINC72008835 | -7.2 | -7.7 | -7.3 | -8.2 | -7.3 | ZINC90644822 | -9.2 | -9.3 | -8.4 | -8.5 | -7.6 |
| ZINC03495890 | -7.9 | -7.2 | -7.1 | -8.1 | -7.4 | ZINC90643013 | -10.1 | -10.1 | -10.2 | -10.2 | -9.2 |
| ZINC14093040 | -8.1 | -8.4 | -9.1 | -8.2 | -7.3 | ZINC33246270 | -8.5 | -9.3 | -7.4 | -9.2 | -7.6 |
| ZINC72366047 | -7.7 | -7.8 | -9 | -8.9 | -10.9 | ZINC33267222 | -9.7 | -7.8 | -9.1 | -9.5 | -7.9 |
| ZINC14000412 | -8.4 | -7.6 | -7.6 | -8.3 | -11.1 | ZINC03434538 | -9.8 | -10.1 | -10.5 | -10.5 | -8.8 |
| ZINC06541302 | -7.8 | -9.1 | -9.3 | -8.3 | -8.1 | ZINC12645453 | -10 | -10.1 | -9.5 | -11.1 | -8.7 |
| ZINC15830923 | -8 | -8.1 | -10.4 | -9.4 | -11 | ZINC15616332 | -8.7 | -8.1 | -7.8 | -8.4 | -7.2 |
| ZINC27008143 | -7.8 | -7 | -7.2 | -8.8 | -8.5 | ZINC17295613 | -10.1 | -8 | -8.7 | -9.4 | -7.3 |
| ZINC60331811 | -8.9 | -8.4 | -9 | -8.6 | -7.9 | ZINC77217043 | -8.9 | -8.8 | -7.5 | -9.6 | -7.8 |
| ZINC08272749 | -8.8 | -10.5 | -9.3 | -8.4 | -7.3 | ZINC19117016 | -9.8 | -8.6 | -9.5 | -8.6 | -8.6 |
| ZINC39916725 | -7.9 | -7.9 | -7.3 | -8 | -11 | ZINC27152565 | -7.9 | -7.6 | -7.8 | -9.2 | -7.6 |
| ZINC16286470 | -7.7 | -8.4 | -7.2 | -8.3 | -7.7 | ZINC19719145 | -8.2 | -7.9 | -8.9 | -8.5 | -7.5 |
| ZINC21727350 | -8.4 | -8.7 | -8.2 | -8 | -7.2 | ZINC83005239 | -9 | -7.8 | -7.9 | -8 | -7.5 |
| ZINC05243732 | -8.3 | -9.9 | -8.3 | -7.6 | -7.1 | ZINC16048649 | -9.3 | -9.6 | -9.2 | -9.5 | -8.8 |
| ZINC44442851 | -7.9 | -7.6 | -7.1 | -8.4 | -7.5 | ZINC04772681 | -9.9 | -9.1 | -9.6 | -10 | -8.8 |
| ZINC27794293 | -8.4 | -8 | -7.9 | -8.1 | -11.8 | ZINC28425593 | -8.5 | -7.6 | -7.4 | -7.8 | -11.9 |
| ZINC77354922 | -8.6 | -7.6 | -9.3 | -7.7 | -10.9 | ZINC30971400 | -8.8 | -8.3 | -8.7 | -8.7 | -11.1 |
| ZINC09874921 | -8.3 | -8 | -8.8 | -7.6 | -11.5 | ZINC23425011 | -8.9 | -7.8 | -8.4 | -8 | -7.8 |
| ZINC09257269 | -8.4 | -8.2 | -8.2 | -8.4 | -7.6 | ZINC77251947 | -9.5 | -8.3 | -8 | -9 | -8.1 |
| ZINC91270382 | -9.1 | -8.7 | -7.7 | -8 | -11.3 | ZINC12340469 | -9.2 | -9.6 | -7.1 | -8.9 | -7.2 |
| ZINC23951108 | -9.3 | -10.3 | -11.6 | -10.4 | -8.2 | ZINC86031007 | -8.7 | -8.3 | -9 | -8.5 | -7.9 |
| ZINC72366044 | -9.1 | -8.5 | -9.8 | -9.1 | -8.7 | ZINC21837772 | -9.5 | -8.8 | -10 | -9 | -8.8 |
| ZINC74161830 | -8.5 | -7.8 | -7 | -8.1 | -13.5 | ZINC44450546 | -9.4 | -8.5 | -8.8 | -8.9 | -8.2 |
| ZINC72366052 | -8.6 | -8.3 | -8.1 | -8.4 | -9.6 | ZINC27706841 | -8.4 | -8.4 | -9.9 | -9.6 | -8.4 |
| ZINC25220594 | -7.1 | -6.4 | -6.3 | -6.3 | -6.5 | ZINC63580102 | -10 | -10.1 | -9.4 | -10.7 | -8.8 |
| ZINC40145195 | -9 | -8.6 | -10.8 | -10 | -12.8 | ZINC68785854 | -10 | -8.4 | -11 | -10.5 | -8.4 |
| ZINC67689452 | -8.8 | -7.3 | -7.8 | -8.1 | -8.2 | ZINC11617357 | -9.4 | -9 | -9.3 | -10 | -7.7 |
| ZINC08709101 | -9 | -10.9 | -10.1 | -9.7 | -7.9 | ZINC23129780 | -9.8 | -9.7 | -11 | -10.3 | -9.1 |
| ZINC95360162 | -8.8 | -9.2 | -8.4 | -8.4 | -10.7 | ZINC81650723 | -10.3 | -8.9 | -10.3 | -9.1 | -9.4 |
| ZINC73308868 | -8.8 | -8.3 | -9.8 | -8.9 | -7.7 | ZINC02659651 | -9.6 | -11.9 | -11.7 | -11 | -9 |
| ZINC04950266 | -8.9 | -7.4 | -9.4 | -8.8 | -12.5 | | | | | | |

Here, METTL1: Methyltransferase-like 1, METTL3: Methyltransferase-like 3, METTL6: Methyltransferase-like 6, METTL16: Methyltransferase-like 16, METTL1: Methyltransferase-like 18.

### S.table 6. Absorption, Digestion, Metabolism, Excretion and Toxicity analysis of top selected drug candidates

| Molecule | ZINC09845394 | ZINC25249143 | ZINC27795763 | ZINC14891501 | ZINC40145195 | ZINC40039977 | ZINC33246308 | ZINC23632079 | ZINC09124297 | ZINC72324946 | ZINC02659651 | CID15516758 1 | CID13540379 8 | CID344265 | CID13903048 7 | CID24858111 |
|---|---|---|---|---|---|---|---|---|---|---|---|---|---|---|---|---|
| **Formula** | C25H16ClN7O3 | C23H19N3O2 | C23H19N3O2 | C20H15ClN2O5 | C25H22N6O | C20H18N4O2S | C25H16ClN3O2 | C18H13N5OS2 | C28H25N5O3 | C21H19N3O3 | C18H12ClN3O3 | C25H28N6O2 | C29H24O12 | C19H14N2O4 | C19H15N5 | C27H23N7O |
| **MW** | 497.89 | 369.42 | 369.42 | 398.8 | 422.48 | 378.45 | 425.87 | 379.46 | 479.53 | 361.39 | 353.76 | 444.53 | 564.49 | 334.33 | 313.36 | 461.52 |
| **#Heavy atoms** | 36 | 28 | 28 | 28 | 32 | 27 | 31 | 26 | 36 | 27 | 25 | 33 | 41 | 25 | 24 | 35 |
| **#Aromatic heavy atoms** | 30 | 23 | 23 | 19 | 24 | 19 | 25 | 22 | 25 | 19 | 19 | 19 | 23 | 15 | 20 | 28 |
| **#Rotatable bonds** | 5 | 4 | 4 | 5 | 4 | 5 | 4 | 3 | 6 | 6 | 4 | 8 | 2 | 4 | 3 | 7 |
| **#H-bond acceptors** | 7 | 3 | 3 | 5 | 4 | 4 | 3 | 4 | 5 | 3 | 4 | 5 | 12 | 4 | 2 | 4 |
| **#H-bond donors** | 1 | 1 | 1 | 1 | 1 | 1 | 2 | 1 | 1 | 3 | 1 | 2 | 9 | 2 | 2 | 4 |
| **MR** | 134.01 | 111.83 | 111.83 | 106.39 | 126.68 | 109.02 | 124.47 | 105.31 | 139.6 | 105.21 | 93.73 | 127 | 143.98 | 94.45 | 94.39 | 140 |
| **TPSA** | 120.73 | 60.05 | 60.05 | 94.45 | 79.18 | 105.12 | 78.61 | 129.48 | 91.04 | 86.98 | 76.61 | 92.8 | 217.6 | 90.47 | 83.42 | 117.85 |
| **iLOGP** | 3.92 | 3.14 | 3.14 | 2.93 | 2.86 | 2.9 | 2.82 | 2.5 | 3.37 | 2.46 | 2.93 | 3.83 | 1.84 | 1.69 | 2.03 | 2.94 |
| **XLOGP3** | 4.18 | 3.32 | 3.32 | 2.66 | 3.29 | 3.08 | 4.72 | 4.44 | 2.94 | 2.38 | 3.03 | 2.72 | 2.38 | 2.61 | 2.92 | 4.78 |
| **WLOGP** | 4.13 | 3.57 | 3.57 | 3.5 | 3.91 | 3.46 | 5.51 | 3.71 | 3.94 | 2.99 | 2.87 | 2.64 | 1.56 | 2.08 | 3.54 | 5.47 |
| **MLOGP** | 2.51 | 3.4 | 3.4 | 2.74 | 3.15 | 2.71 | 3.72 | 3.26 | 3 | 2.12 | 1.26 | 1.93 | -0.79 | 2.32 | 2.02 | 2.39 |
| **Silicos-IT Log P** | 3.02 | 3.35 | 3.35 | 3.96 | 3.35 | 4.5 | 6.76 | 4.2 | 4.1 | 4.58 | 3 | 2.5 | 1.56 | 2.9 | 3.19 | 3.29 |
| **Consensus Log P** | 3.55 | 3.36 | 3.36 | 3.16 | 3.31 | 3.33 | 4.71 | 3.62 | 3.47 | 2.9 | 2.62 | 2.72 | 1.31 | 2.32 | 2.74 | 3.77 |
| **ESOL Log S** | -5.85 | -4.57 | -4.57 | -4.16 | -4.82 | -4.32 | -5.79 | -5.42 | -4.78 | -3.7 | -4.24 | -4.21 | -5.12 | -3.74 | -4.04 | -5.84 |
| **ESOL Solubility (mg/ml)** | 7.08E-04 | 1.00E-02 | 1.00E-02 | 2.76E-02 | 6.35E-03 | 1.82E-02 | 6.96E-04 | 1.45E-03 | 7.90E-03 | 7.13E-02 | 2.03E-02 | 2.76E-02 | 4.26E-03 | 6.12E-02 | 2.85E-02 | 6.63E-04 |
| **ESOL Solubility (mol/l)** | 1.42E-06 | 2.72E-05 | 2.72E-05 | 6.91E-05 | 1.50E-05 | 4.81E-05 | 1.63E-06 | 3.82E-06 | 1.65E-05 | 1.97E-04 | 5.75E-05 | 6.20E-05 | 7.54E-06 | 1.83E-04 | 9.10E-05 | 1.44E-06 |
| **ESOL Class** | Moderately soluble | Moderately soluble | Moderately soluble | Moderately soluble | Moderately soluble | Moderately soluble | Moderately soluble | Moderately soluble | Moderately soluble | Soluble | Moderately soluble | Moderately soluble | Moderately soluble | Soluble | Moderately soluble | Moderately soluble |
| **GI absorption** | High | High | High | High | High | High | High | High | High | High | High | High | Low | High | High | Low |
| **BBB permeant** | No | Yes | Yes | No | No | No | No | No | No | No | Yes | No | No | No | No | No |
| **Pgp substrate** | Yes | Yes | Yes | No | Yes | No | No | No | Yes | Yes | No | Yes | No | No | Yes | No |
| **CYP1A2 inhibitor** | No | No | No | Yes | No | Yes | No | No | No | Yes | Yes | Yes | No | Yes | Yes | Yes |
| **CYP2C19 inhibitor** | No | Yes | Yes | Yes | Yes | Yes | Yes | Yes | Yes | No | No | Yes | No | No | Yes | Yes |
| **CYP2C9 inhibitor** | Yes | Yes | Yes | Yes | Yes | Yes | No | Yes | Yes | Yes | Yes | Yes | Yes | Yes | Yes | No |
| **CYP2D6 inhibitor** | No | Yes | Yes | No | Yes | No | No | No | Yes | Yes | Yes | Yes | No | No | Yes | Yes |
| **CYP3A4 inhibitor** | No | No | No | Yes | Yes | Yes | No | No | Yes | Yes | Yes | Yes | Yes | No | Yes | Yes |
| **log Kp (cm/s)** | -6.37 | -6.2 | -6.2 | -6.84 | -6.54 | -6.42 | -5.55 | -5.46 | -7.14 | -6.81 | -6.31 | -7.08 | -8.05 | -6.49 | -6.14 | -5.72 |
| **Lipinski #violations** | 0 | 0 | 0 | 0 | 0 | 0 | 0 | 0 | 0 | 0 | 0 | 0 | 3 | 0 | 0 | 0 |
| **AMES Toxicity** | No | No | No | No | No | No | No | No | Yes | No | No | Yes | | No | No | No |
| **Carcinogens** | No | No | No | No | No | No | No | No | No | No | No | No | | No | No | No |
| **Biodegradation** | No | No | No | No | No | No | No | No | No | No | No | No | | No | No | No |
| **Acute Oral Toxicity log(1/(mol/kg))** | III, 1.801 | III, 2.347 | III, 2.347 | III 1.741 | III, 2.539 | III 2.742 | III 2.18 | III 1.735 | III, 3.154 | III, 1.983 | III, 1.411 | III 2.492 | | III 1.05 | III 2.668 | III 2.347 |
| **Caco-2 Permeability)** | No | Yes | Yes | Yes | No | No | No | Yes | No | No | No | No | | Yes | No | Yes |
| **Mutagenicity** | No | No | No | No | No | No | No | No | No | No | No | No | No | No | No | Yes |
| **Tumorigenicity** | No | No | No | Yes | No | No | No | No | No | No | No | No | No | No | No | Yes |
| **Irritating effects** | No | No | No | No | No | No | No | Yes | No | No | No | No | No | No | No | No |
| **Reproductive effects** | No | No | No | No | No | No | Yes | No | No | No | No | No | No | Yes | No | Yes |
| **Druglikeness** | 5.74 | 1.59 | 1.59 | -18.69 | 7.18 | 4.22 | 5.13 | 5.77 | 0.09 | 0.74 | 4.89 | -1.68 | 1.67 | -9.13 | -1.4 | -0.55 |
| **Drug-score** | 0.47 | 0.62 | 0.62 | 0.18 | 0.63 | 0.57 | 0.3 | 0.36 | 0.4 | 0.69 | 0.6 | 0.34 | 0.52 | 0.34 | 0.46 | 0.06 |

**S.table 7. Site-specific docking of final drug candidates with different docking tools**

| | | Binding Affinity (Kcal/mol) | | | | |
|---|---|---|---|---|---|---|
| **Drug Candidates** | **Tools** | **METTL1** | **METTL3** | **METTL6** | **METTL16** | **METTL18** |
| ZINC70666503 | Autodock Vina | -9.4 | -10.4 | -11.6 | -9.3 | -13.1 |
| | SwissDock | -8.44 | -8.45 | -8.42 | -7.68 | -7.41 |
| | CB dock | -8.9 | -10.3 | -11.6 | -9.2 | -12.8 |
| ZINC13000658 | Autodock Vina | -9.5 | -10 | -10.0 | -9.6 | -11.8 |
| | SwissDock | -8.38 | -8.67 | -9.24 | -8.63 | -8.67 |
| | CB dock | -9.3 | -9.5 | -10 | -8.9 | -11.7 |
| CID155167581 (Reference) | Autodock Vina | -9.4 | -10.9 | -10.7 | -9.1 | -10.3 |
| | SwissDock | -8.91 | -8.15 | 8.26 | -9.15 | -8.55 |
| | CB dock | -9 | -11.1 | -10.8 | -8.9 | -10.2 |

NOTE: METTL1: Methyltransferase-like 1, METTL3: Methyltransferase-like 3, METTL6: Methyltransferase-like 6, METTL16: Methyltransferase-like 16, METTL18: Methyltransferase-like18

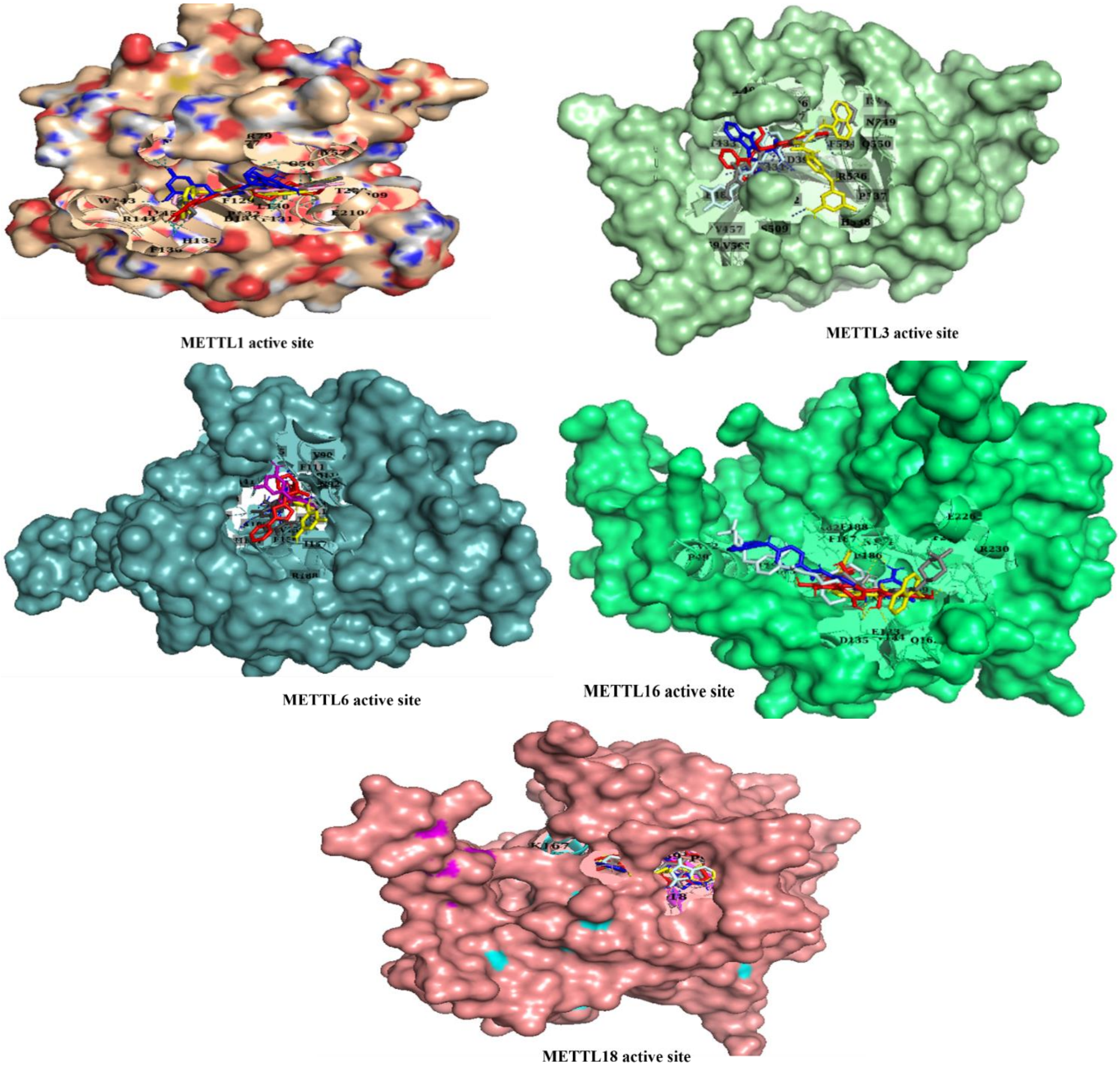


S.figure 1: Active site cavities of METTL1, and METTL3, METTL6, METTL16, and METTL18

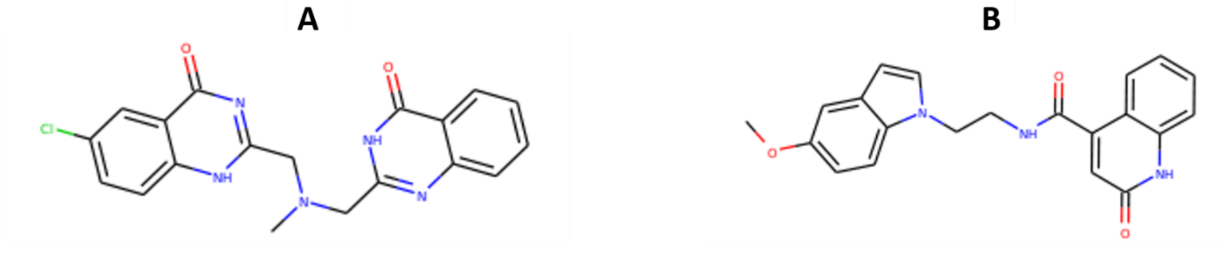


6-chloro-2-({methyl[(4-oxo-3,4-dihydroquinazolin-2-yl)methyl]amino}methyl)-3,4-dihydroquinazolin-4-one

N-(2-(5-methoxy-1H-indol-1-yl)ethyl)-2-oxo-1,2-dihydroquinoline-4-carboxamide

S.figure 2. Chemical structure of ZINC13000658 (AKOS033010880) (A) and ZINC70666503 (EiM08-22770) (B)

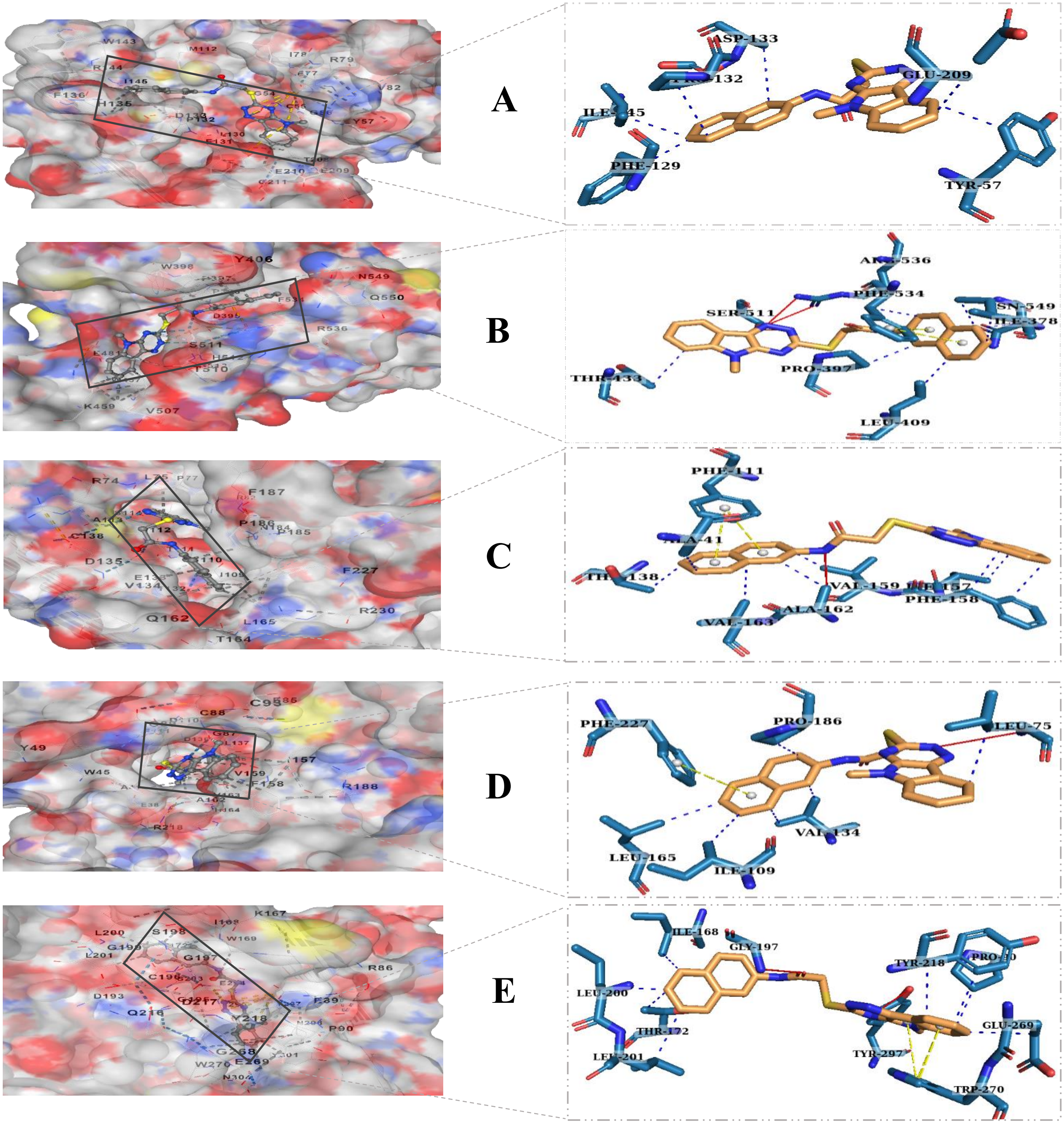


S.figure 3. Binding site analysis of the interaction of ZINC70666503 at METTL1 (A), METTL3 (B), METTL6 (C), METTL16 (D), and METTL18 (E) proteins. The drug-proteins interaction at the protein binding surface (left) is showing largely(right) where Lemon line indicates the drug compound (ZINC70666503) and Sky blue lines indicate interacted amino acids. Here, straight red line is hydrogen bond, and hydrophobic interactions, and π-cation stacking are represented by blue, and yellow dash line, respectively

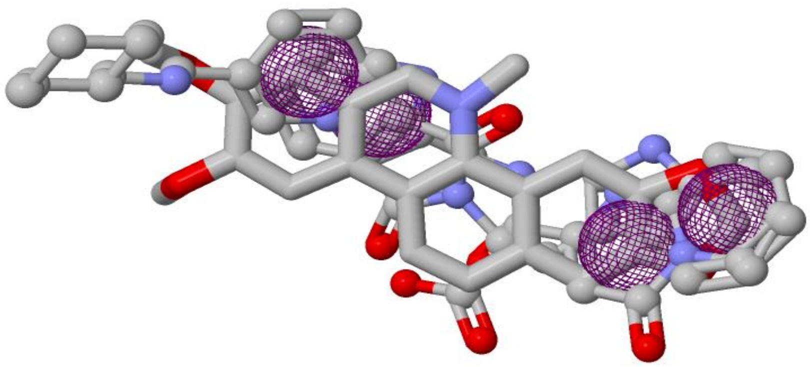

S.figure 4. Ligand-based pharmacophore model. Here, violet represents aromatic features.

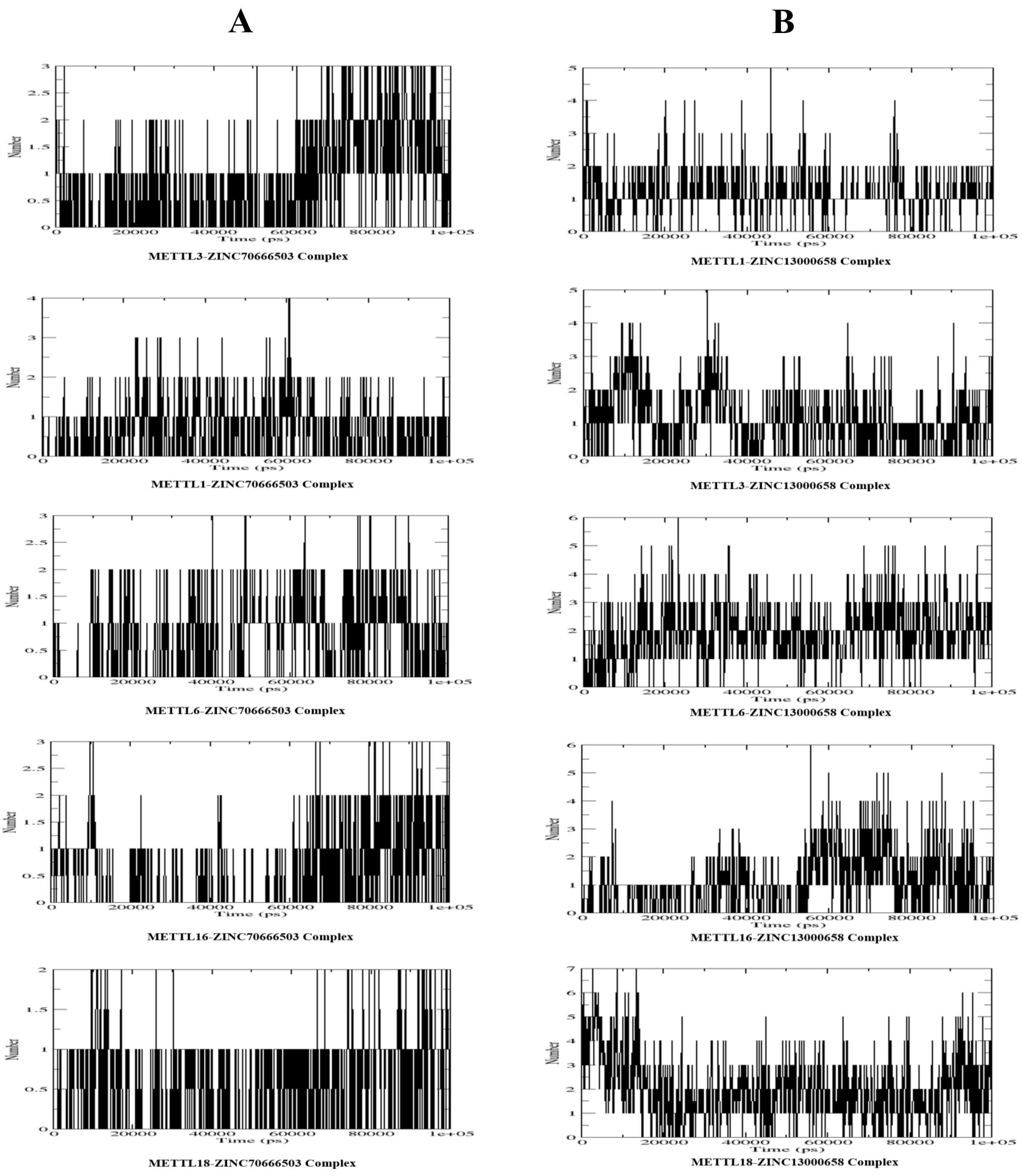


S.figure 5. The statistics of number of hydrogen bonds (H-bonds) involved in the interaction of ZINC70666503 (A) and ZINC13000658 (B) with methyltransferase-like 1,3,6,16, and 18.
